# Cortical cognitive processing explains and predicts superior colliculus signals and timing of gaze shifts to multisensory targets

**DOI:** 10.64898/2026.01.20.700579

**Authors:** Mehdi Daemi, Douglas Crawford

## Abstract

When goal-directed movements are aimed toward multimodal stimuli, action planning differs compared to the unimodal case in both the spatial and temporal domains, as reflected in specific trends of behavioral response differences in the two conditions. This has been systematically reported in past behavioral and neurophysiological studies; however, a unified neurocognitive theory is yet to be proposed that explains all these findings together based on cognitive processes during the delay period.

In our previous paper (Daemi, Harris, & Crawford, 2016) we modeled causal inference in order to determine the *spatial* location of the goal for saccade. Here, we extend this framework into 1) the *temporal* domain, and 2) the domain of neural signals. Specifically, we propose that “confidence” on selecting a winning plan, relative to other alternatives, should influence the timing of execution of the winning action plan, as reflected in known superior colliculus signals and the generated behavior.

To model these concepts, we build upon our previous evidence-accumulation decision-making framework and compute an instantaneous measure of confidence based on the relative saliency of the winning motor plan compared to the alternate plans. A winning plan is only initiated when enough evidence is accumulated in its favor. This is realized by introducing an accumulative measure of confidence that integrates the instantaneous measure through time. A threshold is set on the accumulative confidence and a GO command is released whenever it reaches the threshold. We also formalize the computations of how this model may be neurally implemented in the brain, mainly in the projections between the superior colliculus (SC), the basal ganglia, and the cortex.

This model produces simulations that replicate and explain several experimental multisensory observations. In the behavioral domain, our model shows how higher reaction times are predicted for multi-modal targets with higher reliability, or with higher spatial or temporal disparities, due to less confidence on a unique cause. In the neurophysiological domain, our model replicates the principal multisensory behaviors observed in SC neurons: 1) the dependence of SC neuronal activity on the spatial and temporal structure of cross-modal stimuli (i.e., spatial and temporal principles), 2) the inverse relationship between stimulus intensity and the magnitude of the elicited multisensory response (i.e., principle of inverse effectiveness). We have also simulated some novel predictions that could guide new experimental studies.

Thus, our model provides a unified viewpoint to explain, for the first time, the effects of both spatial and temporal factors on reaction time variability for multisensory targets, and assigns each of these effects to a unique cognitive function upstream from sensorimotor transformations. Our model proposes cognitive significance for multisensory neural principles in SC, by linking them to how the cortex infers a causal structure and calculates confidence on its inference.

## 2 Introduction

We can observe animal behavior. We can record neural activity associated with that behavior. However, we cannot directly see or measure cognitive information processing. To study this, we need theorization. The closest theory to reality is one in which behavioral data is explained, neural data is reproduced, and cognitive architecture is proposed in a unified manner. Here we want to propose such a model for multisensory gaze-shift phenomenon.

### 2.1 Multimodal Reaction Time Behavior

Reaction time (RT) is a measure of speed with which a subject responds to the stimuli within the context of a task. RT has been used to investigate hypotheses about the mental and motor processes to implement different tasks (Sternberg, 1969). In multisensory integration (MSI) research specifically, RT has been used to assess how combining multimodal stimuli with various reliabilities affect task implementation and response generation (Hershenson, 1962; Rubinstein, 1964). Here, we contemplate the possibility of a unified mechanism which can explain the effects of spatial, temporal, and reliability features of cross-modal stimulation on RT.

It is well known that bimodal stimuli, e.g., visual and auditory, affect the reaction times of goal-directed saccadic eye movements. In particular, when the two stimuli are aligned in space and time, a considerable reduction of the saccade RT is typically observed relative to visual stimulus alone or to auditory stimulus alone. Conversely, RT increases more slowly or even decreases when the stimuli are presented farther from each other or when the delay between them gets larger (Corneil, Van Wanrooij, Munoz, & Van Opstal, 2002; Diederich & Colonius, 2004, 2008a, 2008b; Frens, Van Opstal, & Van der Willigen, 1995; Navarra, Hartcher-O’Brien, Piazza, & Spence, 2009; Navarra et al., 2005; Porada, Stein, & Rowland, 2025; Van Wanrooij, Bremen, & John Van Opstal, 2010).

### 2.2 Superior Colliculus Multisensory Neurophysiology

At the level of the single neuron, multisensory integration is defined operationally as: a statistically significant difference between the number of impulses evoked by a cross-modal combination of stimuli and the number evoked by the most effective of these stimuli individually (B. E. Stein, Huneycutt, & Meredith, 1988). To date, the best-studied brain structure for examining multisensory interactions at the neural level has been the superior colliculus (SC) (Corneil, Olivier, & Munoz, 2002; Corneil, Van Wanrooij, et al., 2002; Freedman & Sparks, 1997; Munoz, Dorris, & Klein, 2000; Barry E Stein, Stanford, & Rowland, 2020). SC (Wurtz & Goldberg, 1972), in interaction with the frontal cortex (Schiller, Sandell, & Maunsell, 1987), plays a critical role in the control of both visual fixation and generation of eye-head gaze-shifts (Gandhi & Katnani 2011; Sajad, Sadeh & Crawford 2020).

Collicular saccade neurons can be subdivided into two classes (Munoz & Wurtz, 1995a, 1995b). The burst neurons (BNs) of SGI lack any significant long-lead preparatory activity before saccades, discharge a vigorous burst of action potentials for a restricted range of saccade vectors that defines a closed movement field, and tend to be located in the dorsal part of the intermediate layers. The buildup neurons (BUNs) of SGI have long-lead anticipatory discharges before saccades, discharge for all saccades whose amplitudes are equal to or greater than optimal (i.e., open-ended movement fields), and tend to reside in the ventral part of the intermediate layers (Lee, Helms, Augustine, & Hall, 1997; Meredith & Ramoa, 1998; Munoz & Istvan, 1998). In an awake animal, multisensory integration facilitates the rise to threshold and leads to a reduction in RT, and the neural correlate for this phenomenon is found in the increased pre-target activity of BUNs (Frens, Hepp, Suzuki, & Henn, 1996; Frens & Van Opstal, 1998; Frens, Vanopstal, & Vanderwilligen, 1995; VanOpstal & Frens, 1996). The reduction in monkey RT decreases systematically as the auditory and visual stimuli are moved out of alignment, just as it does for human saccades. In line with this observation, the prelude buildup activity in BUNs is no longer enhanced when the stimuli are misaligned. Instead, they are often depressed. Taken together, multisensory integration is observable in the preparatory phase of saccade programming rather than in the sensory-evoked responses or in the movement-related saccadic burst (Frens & Van Opstal, 1998; VanOpstal & Frens, 1996).

SC neurons’ multisensory behavior has been associated with its afferent connections the most important of which come from cortex (Benedek, Mucke, Norita, Albowitz, & Creutzfeldt, 1988; Clarey & Irvine, 1986; Meredith & Clemo, 1989; Mucke, Norita, Benedek, & Creutzfeldt, 1982). In the lack of cortico-collicular inputs, SC neurons are incapable of integrating cross-modal signals and producing the unique response patterns (Alvarado, Stanford, Vaughan, & Stein, 2007; Jiang, Wallace, Jiang, Vaughan, & Stein, 2001; Wallace, Meredith, & Stein, 1993).

### 2.3 Previous Models

Through the years, there have been various attempts to model the variability of RT in multisensory tasks. They have mostly focused on the effect of temporal configuration of the cross-modal stimuli on the RT. The first group of models is referred to as “separate activation” or “race” models. They assume parallel and completely separate channels of sensory processing for stimuli from different modalities. Each channel builds up some independent activation. Response is triggered by the channel that reaches some threshold level first. Average RT to multisensory stimuli is lower than unimodal stimuli because the average of winner’s processing time is smaller than average processing time in each single channel (statistical facilitation). Independent Gaussian distributions (Raab, 1962) and experimentally observed distributions (Gielen, Schmidt, & Van den Heuvel, 1983) were used as unimodal distributions to estimate the minimum distribution in the bimodal conditions. Nevertheless, statistical facilitation couldn’t account for facilitation in data (Diederich & Colonius, 2008a).

The second group of models is called “coactivation” models. They assume that activation raised in different sensory channels by presenting multimodal stimuli is combined to satisfy a single criterion for response initiation. The discrete realization of this idea gave rise to the so-called superposition models while its continuous realization brought about multichannel diffusion models (Diederich, 1992; Schwarz, 1989). In all such models, the stimulus intensity is represented by some internal indicator (“counter” for superposition models and “drift” for diffusion models). For multimodal stimuli, these internal variables from multiple sensory channels are added together during some peripheral stage of processing. This leads to faster reaching some threshold (fixed number of counts for superposition models and threshold limits for diffusion models) and lower RT.

Previous models could not account for distinguishing a target modality from a non-target modality in experiments like the focused attention paradigm (Amlot, Walker, Driver, & Spence, 2003; Diederich & Colonius, 2007). To consider such effects, time-window-of-integration (TWIN) models combine basic ideas of the previous groups of models (Colonius & Diederich, 2004). They consider two stages of processing. The first stage consists of separate and parallel processing in unisensory pathways. The second stage comprises the combination of the unisensory activations and response initiation. Second stage occurs only if the peripheral processes of the first stage all terminate within a given time interval (Colonius & Diederich, 2010). Such two-stage models support the idea that the race between the sensory channels takes place upstream from the superior colliculus (SC), and that the SC itself is part of the second stage (Sparks & Mays, 1990).

### 2.4 Goals of the Current Study

None of the models, briefly described above, account for effects of spatial disparity of the cross-modal stimuli on the reaction time. These models also ignored the internal perception of the subjects, namely whether they perceive the multimodal stimuli as belonging to a unique event or separate events. Here we propose a model that explains the variability of saccadic reaction time as a function of both temporal and spatial configurations of the multimodal stimuli. We build this model upon a previous model of causal inference for spatial localization in multi-modal situations (Daemi et al., 2016). Before, a decision-making circuitry was proposed, where different plans represented different possibilities for the inferred cause, namely same or separate sources. A spatiotemporal similarity measure was introduced, and was compared to the reliabilities of the unimodal stimuli to make the decision on the causal structure.

Now, we extend that framework to model how the timing of an action based on the inference is determined. The saccadic RT is proposed to depend on a measure of accumulated confidence on the decision made about the causal structure, as seen before in parietal neurons (Kiani & Shadlen, 2009; Vivar-Lazo & Fetsch, 2025). This expanded framework can explain variability of the RT as functions of 1) spatial configuration of the stimuli 2) temporal configuration of stimuli 3) reliabilities of the stimuli. This means we can explain the interesting patterns of variability in reaction times of gaze-shifts towards cross-modal stimuli, which could not be explained based on psychophysical models of sensory-driven reactions. We do so by proposing an internal model and assigning a cognitive significance, namely accumulative confidence, as the factor governing the RT.

In the realm of the neurophysiology of multisensory gaze shifts, the big question is how we can explain these various neurophysiological phenomena, mentioned above, in a common framework and in relation to sensorimotor, and possibly cognitive, gaze-shift planning. To address this question, taking the approach of the neural engineering framework (Dumont et al., 2023; C. Eliasmith et al., 2012), which unifies the behavioral and neurophysiological aspects, we neurally implement the causal inference (Daemi et al., 2016) and reaction time (first part of this paper) models. We propose a model of the functional interactions between SC, Basal Ganglia, and the cortical decision-making circuitry underlying planning a gaze-shift. Our model relates the behavioral level to the underlying neurophysiology in a biologically plausible manner. It replicates the known neural behavior of SC and explains it in relation to causal inference and action planning. It explains the effect of cortico-collicular projections on SC multisensory behavior in relation to cognitive phenomena.

## 3 Model Overview

This section provides a verbal/conceptual description of our model (Figure 1). For a mathematical description, see the Appendix below.

**Figure 1:**
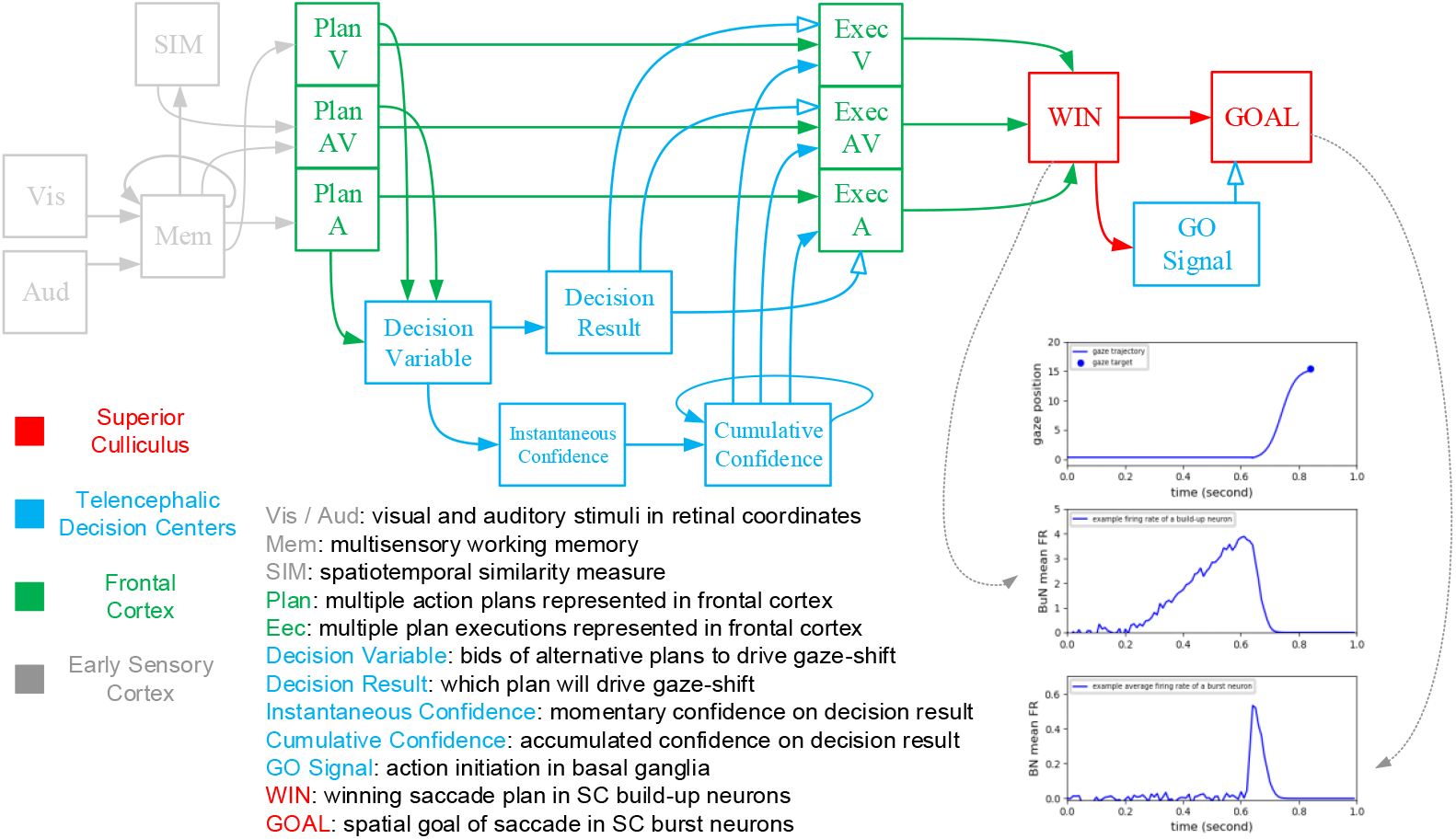
The model of reaction time variability of gaze-shifts towards cross-modal stimuli. Upstream of this model, in early unisensory cortex (grey), the spatiotemporal similarity between visual and auditory stimuli is measured and the multimodal signals were integrated in working memory. The three possible plans are constructed in units 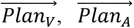, and 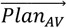 as saliency spatial maps. The decision variable is constructed in the unit 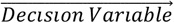 by sending the saliency of the three plans as their bids. The decision is made in the decision result 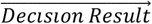, by materializing the concept that if the similarity measure is bigger than a threshold, then the multisensory plan wins and if not, the unisensory plan with higher reliability wins. The 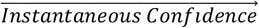, and 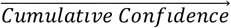 on the result of the decision are calculated from the 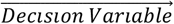. The plan representations in the execution layer, three units 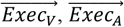, and 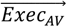, are constructed as confidence maps by communicating the spatial information from plan layer and the corresponding accumulative confidence value. The decision result is realized by selective inhibition of the plan units in execution layer (not shown). The confidence map of the winning plan is then sent to the unit 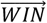. The confidence value of the winning plan is sent to 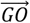. The spatial information of the 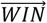 is sent to 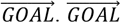 is under constant inhibition of 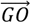. When the confidence reaches a threshold, 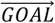 is disinhibited.

Generally, in terms of action initiation, two types of tasks are possible: 1) a forced-reaction-time (forced-RT) task, and 2) a choice-reaction-time (choice-RT) task. In the forced-RT task the time for onset of the action is a requirement of the task and is forced by a higher-order, top-down command. In the choice-RT task the subject is free to start the action as soon as it is ready. Here, we want to build a model of gaze-shift initiation, in multi-modal situations, for a choice-RT task. We propose that the readiness for initiating a winning action plan is measured by the confidence on the decision that determined that plan is winning.

Consider a situation where visual and / or auditory stimuli, with possible spatial and temporal disparities and different reliabilities are presented to a subject who is instructed to make a gaze-shift to the most reliable of the targets. We previously proposed a model of how subjects may infer the cause of the stimuli, a common source or separate sources, by introducing a spatiotemporal measure of similarity (Daemi et al., 2016). Here we extend that model to account for the variability of multi-modal reaction times. Figure 1 illustrates this extended, unifying framework, containing three color-coded sections, explained in the following paragraphs.

The green parts of the model include the decision-making circuitry underlying causal inference. This is where the three alternative plans, realizing the three possible solutions to the causal inference problem, are represented. These include a unisensory plan 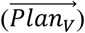 that manifests separate sources and gaze-shift to the visual signal, another unisensory plan 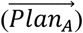 that manifests separate sources and gaze-shift to the auditory signal, and a multisensory plan 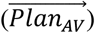 that manifests a common source and gaze-shift to the weighted average of their spatial positions. Each of these plans include a saliency component, and the decision on the causal structure is made by systematically comparing these saliencies. The saliencies of the unisensory plans are the reliabilities of the unimodal stimuli (that can be simplified to their intensities). The saliency of the multisensory plan is a measure of spatiotemporal similarity between the cross-modal stimuli. For making the decision, the saliencies of the alternative plans are sent to construct a decision variable 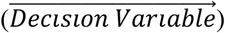. A decision rule is then applied on the decision variable in its transformation to the decision result 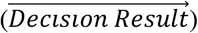. The decision rule realizes a specific comparison between the plan saliencies: the multisensory plan is chosen if its saliency is greater than a threshold, and the more reliable of the unisensory plans is chosen if the similarity measure is smaller than the threshold.

The blue parts of the model are involved in calculating *when* to send the winning plan for execution and initiation of the action. An instantaneous measure of confidence 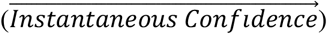 is first constructed by transforming the decision variable, realizing the idea of immediate confidence on a plan to guide action if it is winning the competition among alternative plans. So, at any time, the component corresponding to a plan is zero, if that plan is not winning, and is equal to the momentary confidence on the decision that the plan is winning, if it is winning. As the decision result could be changing by accumulation of evidence, a component could be changing between zero and a positive value through time. Then, an accumulative measure of confidence 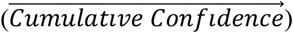 is constructed by integration, through time, of the instantaneous measure of confidence. This measure reflects accumulation of evidence, through time, against or in favor of a plan to guide the action, as conceptualized before in other studies (Kiani & Shadlen, 2009). Each component indicates the amount of confidence on implementing a plan, accumulated through time.

We have salience maps of space in the plan layer (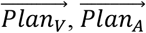, and 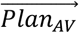), which represented alternative causal structure. In order to implement the decision on the preferred causal structure, another layer of plan representations is constructed in an execution layer. This involves three confidence maps of space, which receive their spatial components from the corresponding representations in the plan layer, and their confidence components from corresponding components of the accumulative measure of confidence. They include a unisensory map 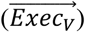 that manifests the victory of the visual plan, another unisensory map 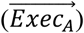 that manifests the victory of the auditory plan, and a multisensory map 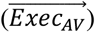 that manifests the victory of the multisensory plan. These plan representations are selectively inhibited by the decision result to implement the selection of the inferred causal structure.

The red parts of the model are involved in timing of action initiation. In previous parts, a spatial plan was chosen to guide the action, and the confidence on the decision to choose that plan was calculated. All this information is reflected in the one confidence map of space, in the execution layer, which was disinhibited by the decision result. That winning plan is now communicated to a computational unit called 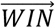. The spatial plan, only, is then communicated from 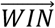to another computational unit called 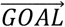. However, 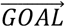 is constantly inhibited by a 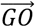command, by default, when we are not confident enough to execute a plan. The 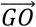command is constructed by applying a threshold function on the confidence measure in 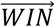. The result of this function is that the 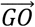 signal is ‘zero’ if the 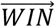 ‘s confidence measure is smaller than a threshold and it becomes ‘one’ if the confidence rises above the threshold. 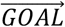 is inhibited by 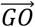 if 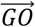 is ‘zero’. However, if 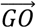 becomes ‘one’ the 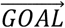 is disinhibited and the spatial plan of WIN is allowed to be communicated into 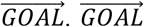 then sends the winning plan to the machinery involved in eye-head coordination and implementing the gaze-shift.

The general outline of the model and the structure proposed in it is inspired by known properties of the decision making, action selection, and gaze control systems in the brain. The model’s representations of alternative plans involved in decision making, i.e. the plan and execution layers, are inspired by such neural codes in frontal cortex (Berendse, Galis-de Graaf, & Groenewegen, 1992; Canteras, Shammah-Lagnado, Silva, & Ricardo, 1990; Jones, Coulter, Burton, & Porter, 1977; Levesque, Charara, Gagnon, Parent, & Deschenes, 1996; Yeterian & Pandya, 1994). A central arbitrating system is thought to receive bids, namely plan saliencies as in our case, from alternative plans for further processing (Redgrave, Prescott, & Gurney, 1999). This information processing, e.g., in the telencephalic decision centers, underlies constructing a decision variable, implementing a decision rule, computing a decision result, and calculating the confidence on the decision. The decision making units and the internal connections between them and their projections from plan representations have been inspired by the known physiology of ubiquitous decision making system in the brain (Cisek & Kalaska, 2010; Gold & Shadlen, 2007). The basal ganglia are thought to receive the result of the decision from cortex (Beiser & Houk, 1998; Gernert, Hamann, Bennay, Loscher, & Richter, 2000; Koos & Tepper, 1999) and implement it through selective disinhibition of cortical and sub-cortical representations. This is realized in our model, through a multiplicative effect on plan representations, in two occasions: 1) selective inhibition of the execution layer by the 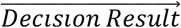, 2) selective inhibition of the goal representation 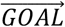 by the ‘go’ command 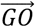. We assume the winning plan in execution layer is sent to the gaze control system to plan a gaze-shift (while it could possibly be sent to other motor circuitries to plan a reach or grasp, for example). So, our final spatial maps are specifically inspired by the saccade-related neural populations in the superior colliculus (Munoz & Wurtz, 1995c, 1995d): 1) confidence map in 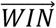 is thought to be implementing the function of the buildup neurons, as previously suggested (Ratcliff, Cherian, & Segraves, 2003; Ratcliff, Hasegawa, Hasegawa, Smith, & Segraves, 2007), although in isolation. 2) motor map in 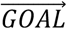 is inspired by the physiology of the burst neurons. The final winning plan 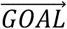 is then assumed to be sent to the brainstem (Girard & Berthoz, 2005; Sparks, 2002) to drive the eye-head coordination system (Daemi & Crawford, 2015; Klier, Wang, & Crawford, 2003) to reorient the line of sight to the selected target.

## 4 Method

The proposed model incorporates a concept of “confidence” on the selected plan, within a decision-making framework, as the underlying factor that determines the timing of plan’s execution (see Model Overview). We show that this concept can explain the complex variability of reaction times in cross-modal configurations (see Results). We are linking the time dimension of action planning to the dynamics of evidence-based decision making, which is a high-level cognitive process. To model this, we need to move 1) beyond the classic cognitive architectures that neglect the time dimension of inferential and logical transformations (Anderson, 1983; Newell & Simon, 1972), and 2) beyond the traditional approaches, like classic-control theory and cybernetics, which constrain goal-directed motor planning with real-time constraints of environmental interactions, but ignore the high-level cognitions (van Gelder, 1998).

A more general framework realizes cognitive processes within the time constraints of interacting with and surviving in a dynamically changing environment, just like the brain. It considers “perception-action” and “high-level cognition” in a unified framework (Chris Eliasmith, 2013). Inspired by the brain neurophysiology (Joaquin M. Fuster, 2005), models within this more general framework are implemented in distributed networks of parallel processing units. This characterizes sensorimotor and cognitive transformations by functions of both the internal state variables and the time, realized in connections between the units. Routing of information between the units (attentional control), through time, is flexibly controlled.

We used the Nengo software (Bekolay et al., 2014) to implement the model, test it through various simulations, and gather behavioral and neural data. Nengo is a Python package for building, testing, and deploying spiking neural networks. We ran the simulations on a personal computer with an i5-3317U CPU @ 1.7GHz. We used Visual Studio 2019 as Python IDE. We used Python packages Matplotlib for visualizations and NumPy and SciPy for further data analysis.

## 5 Results

Here we use our model to simulate four previous and three novel experimental paradigms. We first verify the internal mechanisms suggested in the model by reproducing the wide range of the variability of RT and SC behavior during previously tested tasks, and then make predictions for some novel non-tested situations. In each task, a specific sensory feature of the gaze-shift target is varied: 1) reliability of stationary unisensory stimulus, 2) spatial distance between stationary cross-modal stimuli, 3) temporal offset between stationary cross-modal stimuli, 4) reliability of stationary cross-modal stimuli, 5) speed of distancing linear movement of one of the two cross-modal stimuli, 6) amplitude of vibrating movement of one of the two cross-modal stimuli, 7) spatial distance between flickering cross-modal stimuli. Then, we will summarize the behavioral results in the last section where we draw the reaction time as functions the changing feature.

### 5.1 Classic Experimental Paradigms

Here, we have simulated previous experimental studies and reproduced their reported trends in results, in order to verify that our model is a good model.

#### Unisensory condition

It has been experimentally observed that when making a gaze-shift towards unimodal stimuli, the reaction time and response onset latency of SC burst neurons decrease, when the intensity of the stimulus increases (Bell, Meredith, Van Opstal, & Munoz, 2006). There have been previous models that were explicit about accumulating unimodal salience signals as evidence for or against some decision alternative (Purcell et al., 2010; Purcell, Schall, Logan, & Palmeri, 2012). Here we simulate this task with a single visual target (Fig 2), assuming reliability is the same as intensity for a stationary dot stimulus. The reliability of the visual stimulus changes in four levels (Fig 2A), while spatial and temporal features stay the same. RT decreases when the unisensory stimulus reliability increases (Fig 2B), reflected in the faster build-up of activity in BuNs (Fig 2C) and earlier response onset of BNs (Fig 2D). This is accomplished in the model by implementing the idea that when there is no distracting stimulus, the confidence (Fig 2F, 2G) on a unique target for shifting the attention increases when its reliability increases (Fig 2E), and the Go signal is issued faster (Fig 2H).

**Figure 2:**
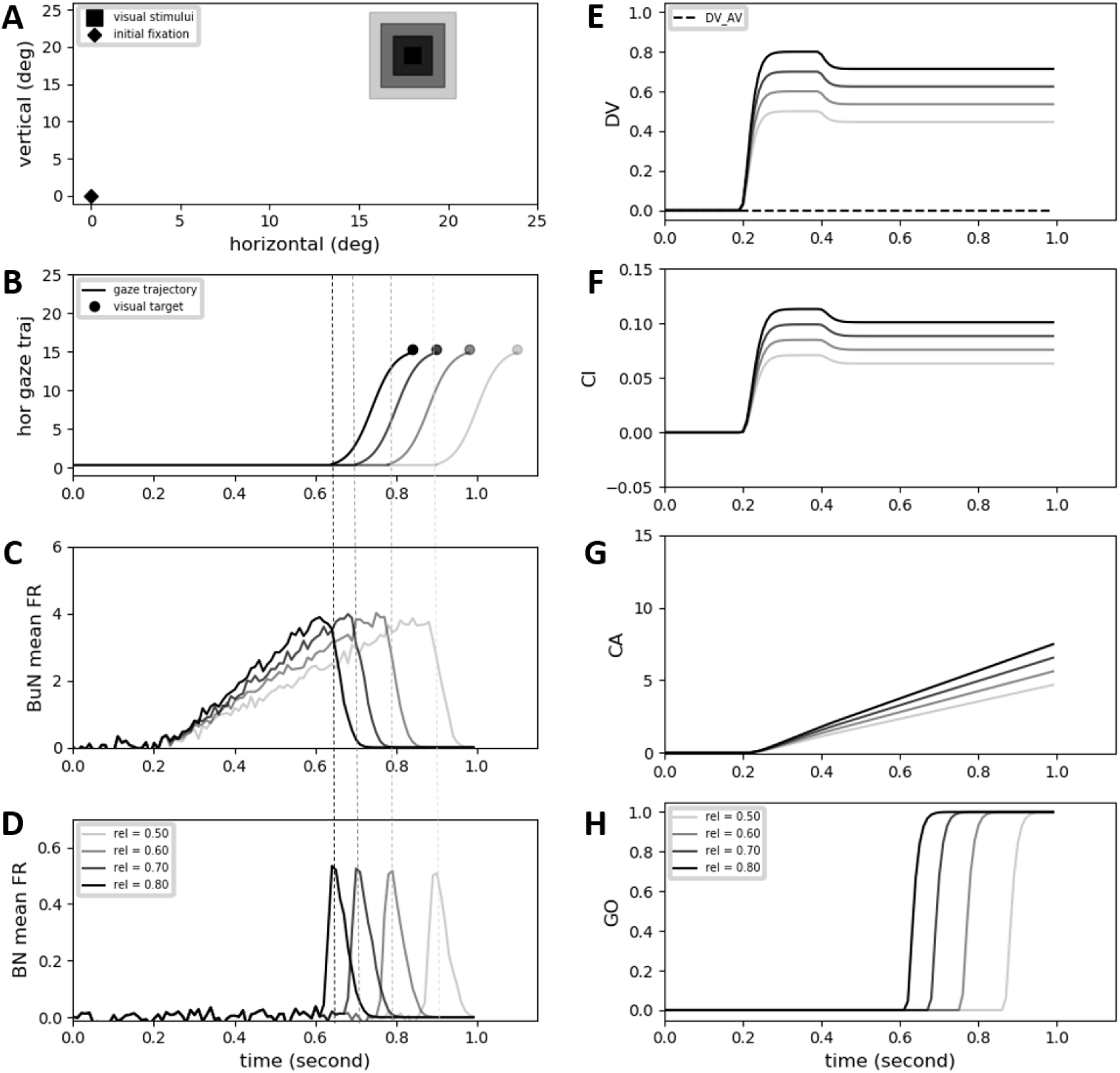
Effect of unimodal target reliability on reaction time. The auditory target is not presented. The position, onset time, and duration of the visual target is fixed. **A)** The reliability of the visual target varies within four different conditions, as illustrated by different levels of blurriness. **B)** Gaze position is shown as a function of time. It shows the reaction time in various conditions. **C)** Mean firing rate of build-up neurons in superior colliculus (SC) is shown as a function of time. **D)** Mean firing rate of burst-neurons in SC is shown as a function of time. **E)** The decision variable is shown being developed through time. The multisensory and auditory components are zero for all conditions. The visual component is changing between conditions because the reliability (rel) of visual target varies. **F)** The instantaneous confidence on decision is shown through time. Only the visual component has non-zero value. **G)** The accumulative confidence on decision is shown as a function of time. Only the visual component is not zero. **H)** The GO signal is shown for execution of the winning plan. It changes from zero to one whenever the accumulative confidence passes a threshold.

#### Effect of spatial disparity

Reaction time of gaze-shifts towards cross-modal stimuli increases, along with the increase in response onset latency of the SC burst neurons, if the spatial distance between the two presented stimuli increases, as experimentally observed (Frens, Van Opstal, et al., 1995; Meredith & Stein, 1986). Here we simulate this task (Fig 3) with visual and auditory stimuli having the same fixed temporal features (onset time 0.2s and duration 0.3s). The position of the visual stimulus is invariable while the position of the auditory stimulus is changing within five conditions, from 0.75° to 8.62° from the visual stimulus (Fig 3A). RT increases when the spatial distance increases (Fig 3B), reflected in the slower build-up of activity in BuNs (Fig 3C) and later response onset of BNs (Fig 3D). This is accomplished in the model by implementing the idea the confidence (Fig 3F, 3G) on the sameness of the origin of the stimuli decreases when the spatial distance between the stimuli increases (Fig 3E), and the Go signal is issued slower (Fig 3H).

**Figure 3:**
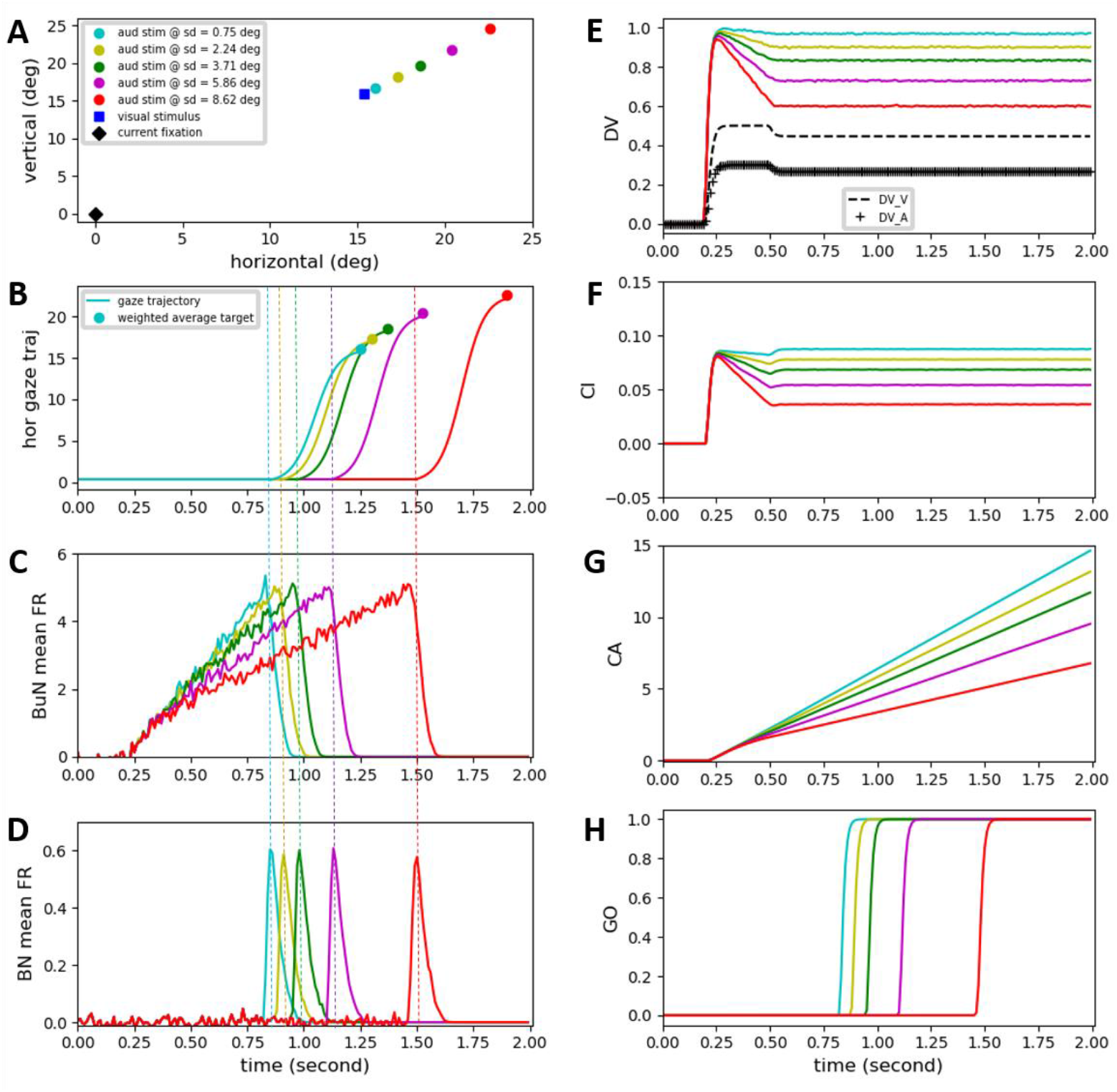
Effect of spatial disparity of cross-modal stimuli on reaction time. Five different conditions have been considered (illustrated by color coding). The visual stimulus has a fixed position, onset time, duration, and reliability for all conditions. The onset time, duration and reliability of the auditory target is also fixed. **A)** The spatial position of the auditory target varies, which is reflected in different spatial distances (sd) for different conditions, while the visual target is fixed. **B)** Gaze position is shown as a function of time. It shows the reaction time in various conditions. **C)** Mean firing rate of build-up neurons in superior colliculus (SC) is shown as a function of time. **D)** Mean firing rate of burst-neurons in SC is shown as a function of time. **E)** The decision variable is shown being developed through time. The unimodal components are the same for all conditions (dashed and crossed lines). The multisensory component is changing between conditions because the spatial distance varies. **F)** The instantaneous confidence on decision is shown through time. The unimodal components are always zero because the multisensory plan is always winning, because its saliency, i.e. the similarity measure, is greater than the threshold. **G)** The accumulative confidence on decision is shown as a function of time. Only the multisensory component is not zero. **H)** The GO signal is shown for execution of the winning plan. It changes from zero to one whenever the accumulative confidence passes a threshold.

#### Effect of temporal disparity

As experimentally observed depressions (Frens, Van Opstal, et al., 1995; Meredith, Nemitz, & Stein, 1987), the reaction time of gaze-shifts towards cross-modal stimuli increases, along with the increase in response onset latency of the SC burst neurons, if the temporal disparity between the two presented stimuli increases. Here we simulate this task (Fig 4) where the visual and auditory stimuli are presented at the same positions and with the same time duration (0.3 (s)) all the time. While the onset time of the visual target is invariable (0.2 (s)), the onset time of the auditory target changes within five conditions from 0.215 (s) to 0.275 (s) (Fig 4A). RT increases when the temporal offset increases (Fig 4B), reflected in the slower build-up of activity in BuNs (Fig 4C) and later response onset of BNs (Fig. 4D). This is accomplished in the model by implementing the idea that when there is no distracting stimulus, the confidence (Figs 4F, 4G) on the unique-object causal structure decreases when the temporal distance between the stimuli increases (Fig 4E), and the Go signal is issued slower (Fig 4H).

**Figure 4:**
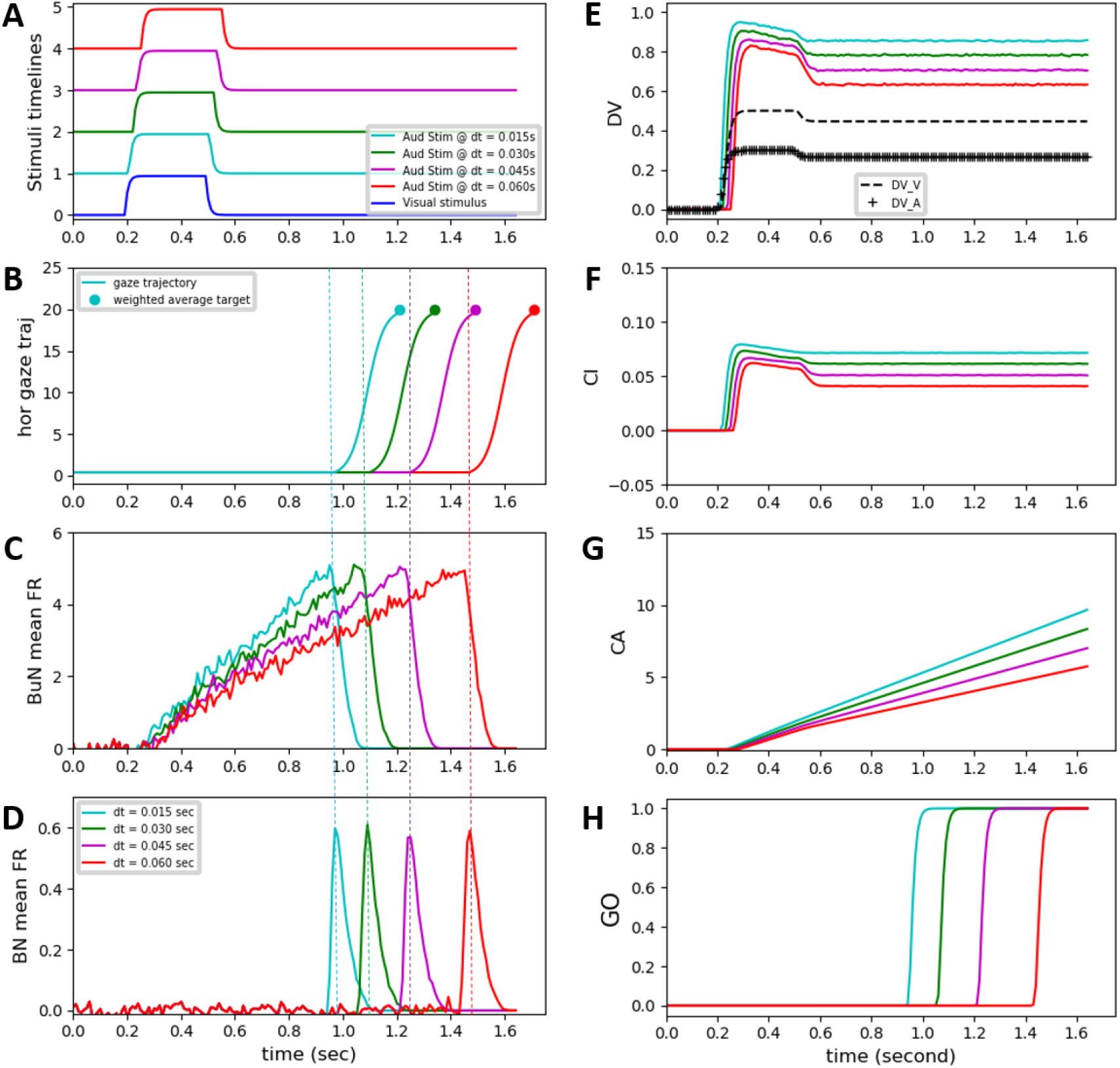
Effect of temporal disparity of cross-modal stimuli on reaction time. Five different conditions have been considered (illustrated by color coding). The visual target has a fixed position, onset time, duration, and reliability for all conditions. The position, duration and reliability of the auditory target is also fixed. **A)** The onset time of the auditory target varies changes relative to the onset time of the visual target from temporal disparity (dt) of 0.015 to 0.075 (s). **B)** Gaze position is shown as a function of time. It shows the reaction time in various conditions. **C)** Mean firing rate of build-up neurons in superior colliculus (SC) is shown as a function of time. **D)** Mean firing rate of burst-neurons in SC is shown as a function of time. **E)** The decision variable is shown being developed through time. The unimodal components are the same for all conditions. The multisensory component is changing between conditions because the spatial distance varies. **F)** The instantaneous confidence on decision is shown through time. The unimodal components are always zero because the multisensory plan is always winning, because its saliency, i.e. the similarity measure, is greater than the threshold. **G)** The accumulative confidence on decision is shown as a function of time. Only the multisensory component is not zero. **H)** The GO signal is shown for execution of the winning plan. It changes from zero to one whenever the accumulative confidence passes a threshold.

#### Inverse effectiveness

In section 4.1 we showed that for a unimodal stimulus, the reaction time decreases by increasing the reliability of the stimulus, as seen in experiments (Bell et al., 2006). However, it has been experimentally seen (Diederich & Colonius, 2004) that, for the multisensory case, this effect is reversed and the reaction time increases by increasing the reliability of the cross-modal stimuli. Here we simulate this task with a single visual target (Fig 5), where visual and auditory stimuli are presented at fixed positions and with invariant temporal features all the time. However, the reliabilities of the two stimuli are the same but changing within four conditions (Fig 5A). RT increases when the multisensory stimulus reliability increases (Fig 5B), reflected in the slower build-up of activity in BuNs (Fig 5C) and later response onset of BNs (Fig 5D). This is accomplished in the model by implementing the idea that the confidence (Figs 5F, 5G) on the sameness of the source of the cross-modal signals decreases if the reliability of the unimodal components increases (Fig 5E), and the Go signal is issued slower (Fig 5H).

**Figure 5:**
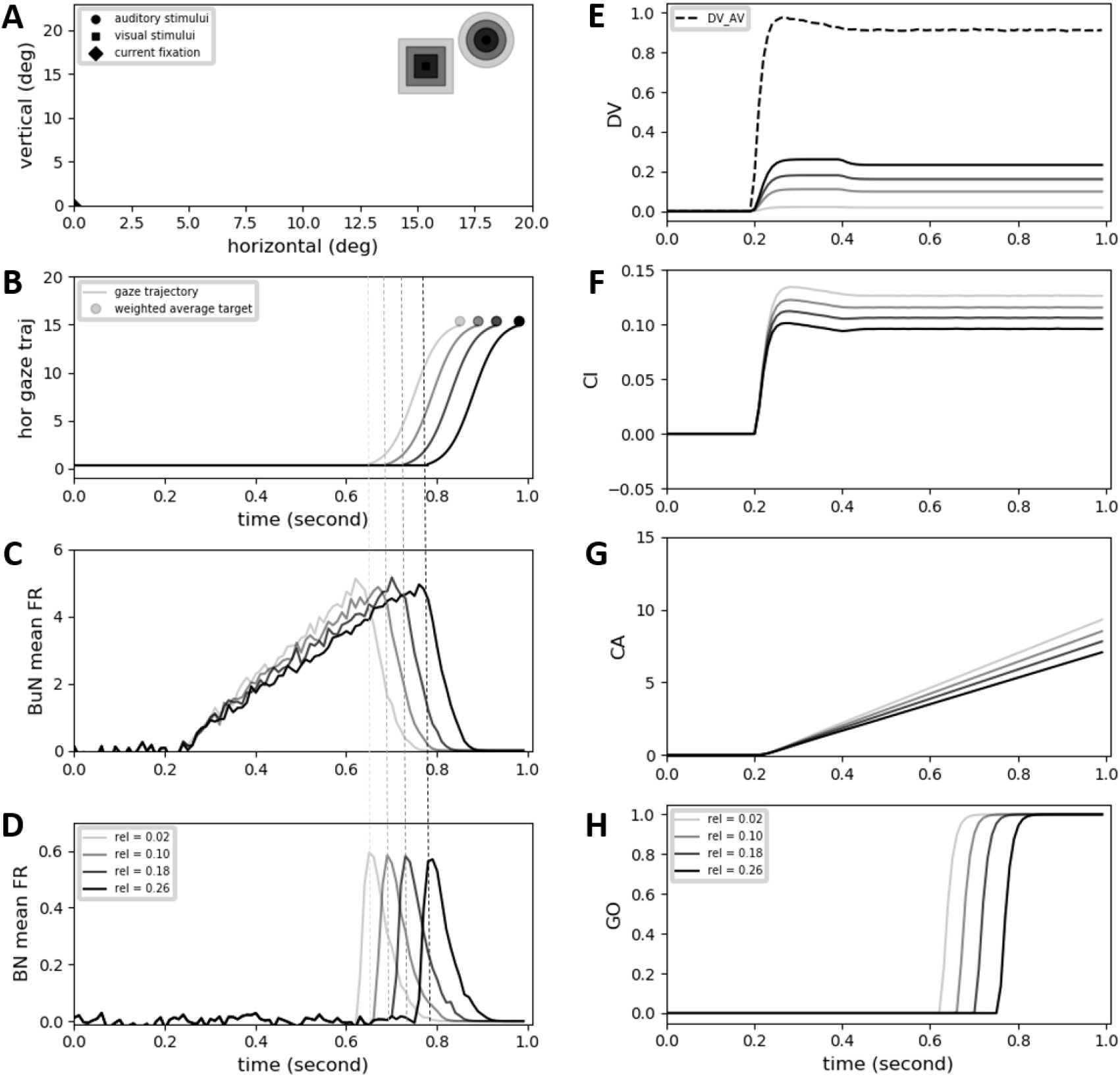
Effect of the reliability of cross-modal stimuli on reaction time. Four different conditions have been considered for all of which both visual and auditory targets have fixed positions, onset times, and durations. **A)** The reliability of the stimuli are the same for the two modalities but changing between different conditions, as illustrated by the varying levels of blurriness of stimuli. **B)** Gaze position is shown as a function of time. It shows the reaction time in various conditions. **C)** Mean firing rate of build-up neurons in superior colliculus (SC) is shown as a function of time. **D)** Mean firing rate of burst-neurons in SC is shown as a function of time. **E)** The decision variable is shown being developed through time. The multisensory component is the same for all conditions. The unimodal components are changing between conditions (while they are the same for the two modalities in one condition) because the reliabilities vary. **F)** The instantaneous confidence on decision is shown through time. The unimodal components are always zero because the multisensory plan is always winning, because its saliency, i.e. the similarity measure, is greater than the threshold. **G)** The accumulative confidence on decision is shown as a function of time. Only the multisensory component is not zero. **H)** The GO signal is shown for execution of the winning plan. It changes from zero to one whenever the accumulative confidence passes a threshold.

### 5.2 Novel Experimental Paradigms

Here, we have come up with novel experimental paradigms, which have not been tested before. This shows how our model can extend our understanding of multisensory animal behavior and the underlying neurocognitive processes.

#### Effect of linear, radial, movement

A non-tested cross-modal gaze-shift task is when one of the stimuli is moving and gradually distancing from the other. Here we simulate this task (Fig 6), assuming the same onset, duration, and offset for the both stimuli. The visual stimulus is stationary but auditory stimulus is moving with constant speed further. The movement speed of the auditory stimulus changes in four levels (Fig 6A). RT increases when distancing movement speed increases (Fig 6B), reflected in the slower build-up of activity in BuNs (Fig 6C) and later response onset of BNs (Fig 6D). This implements the idea that the confidence (Fig 6F, 6G) on the uniqueness of the origin of the cross-modal stimuli decreases when one of them is moving with higher speed (Fig 6E), so, the Go signal is issued slower (Fig 6H).

**Figure 6:**
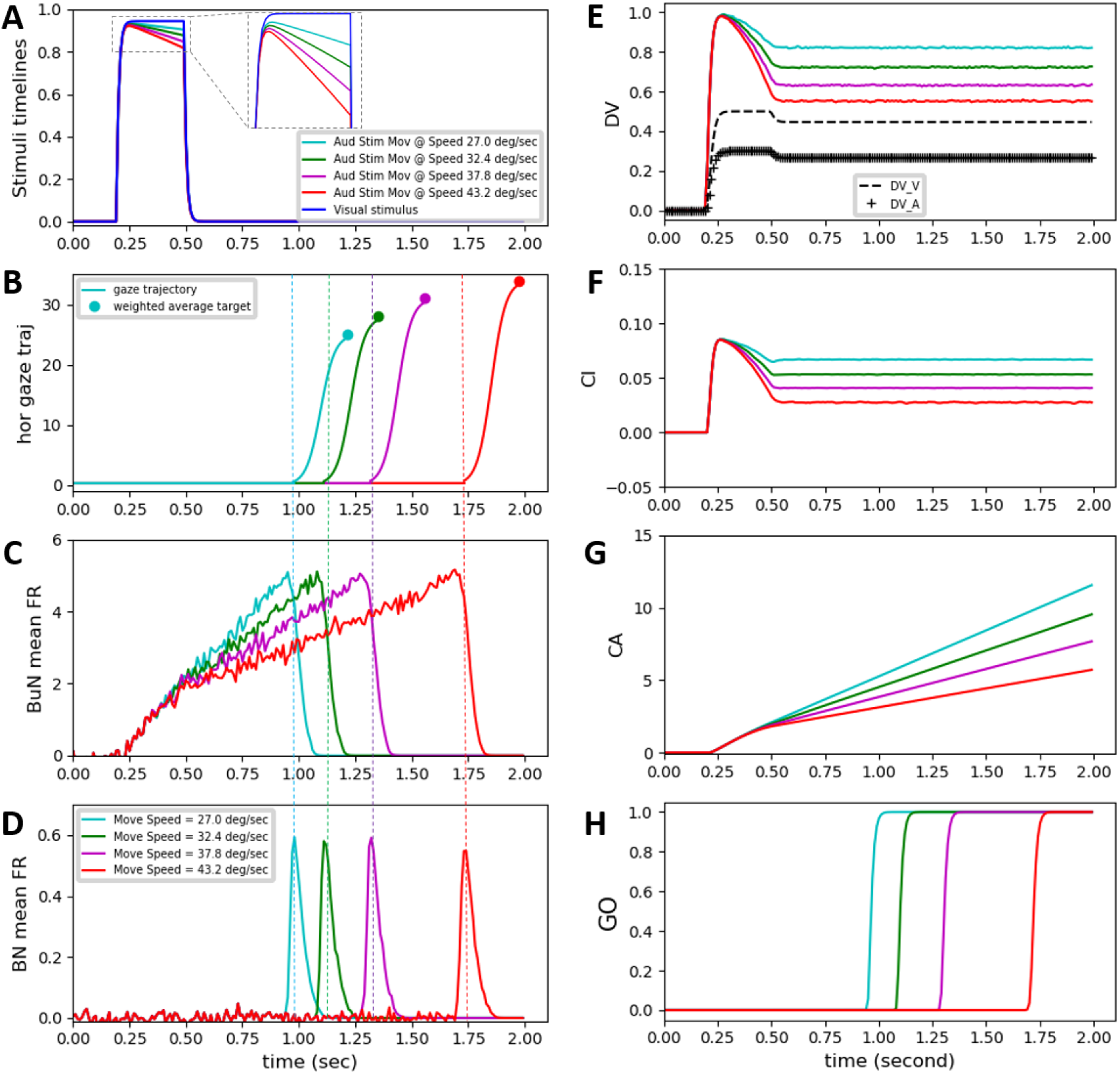
Effect of linear movement of one stimulus on reaction time. Four different conditions have been considered (illustrated by color coding). The visual stimulus has a fixed position, onset time, duration, and reliability for all conditions. The onset time, duration and reliability of the auditory target is also fixed. **A)** The auditory stimulus moves with different speeds further from the stationary visual stimulus. **B)** Gaze position is shown as a function of time. It shows the reaction time in various conditions. **C)** Mean firing rate of build-up neurons in superior colliculus (SC) is shown as a function of time. **D)** Mean firing rate of burst-neurons in SC is shown as a function of time. **E)** The decision variable is shown being developed through time. The unimodal components are the same for all conditions (dashed and crossed lines). The multisensory component is changing between conditions. **F)** The instantaneous confidence on decision is shown through time. The unimodal components are always zero because the multisensory plan is always winning. **G)** The accumulative confidence on decision is shown as a function of time. Only the multisensory component is not zero. **H)** The GO signal is shown for execution of the winning plan. It changes from zero to one whenever the accumulative confidence passes a threshold.

#### Effect of vibrating movement

A second non-tested cross-modal gaze-shift task is when one of the stimuli is vibrating in space, back and forth, around the other fixed one. Here we simulate this task (Fig 7), assuming the same onset, duration, and offset for the both stimuli. The visual stimulus is stationary but auditory stimulus is going back and forth. The amplitude of vibration of the auditory stimulus changes in four levels (Fig 7A). RT increases when vibration amplitude increases (Fig 7B), reflected in the slower build-up of activity in BuNs (Fig 7C) and later response onset of BNs (Fig 7D). This implements the idea that the confidence (Fig 7F, 7G) on the uniqueness of the origin of the cross-modal stimuli decreases when one of them is moving back and forth with higher amplitude (Fig 7E), so, the Go signal is issued slower (Fig 7H).

**Figure 7:**
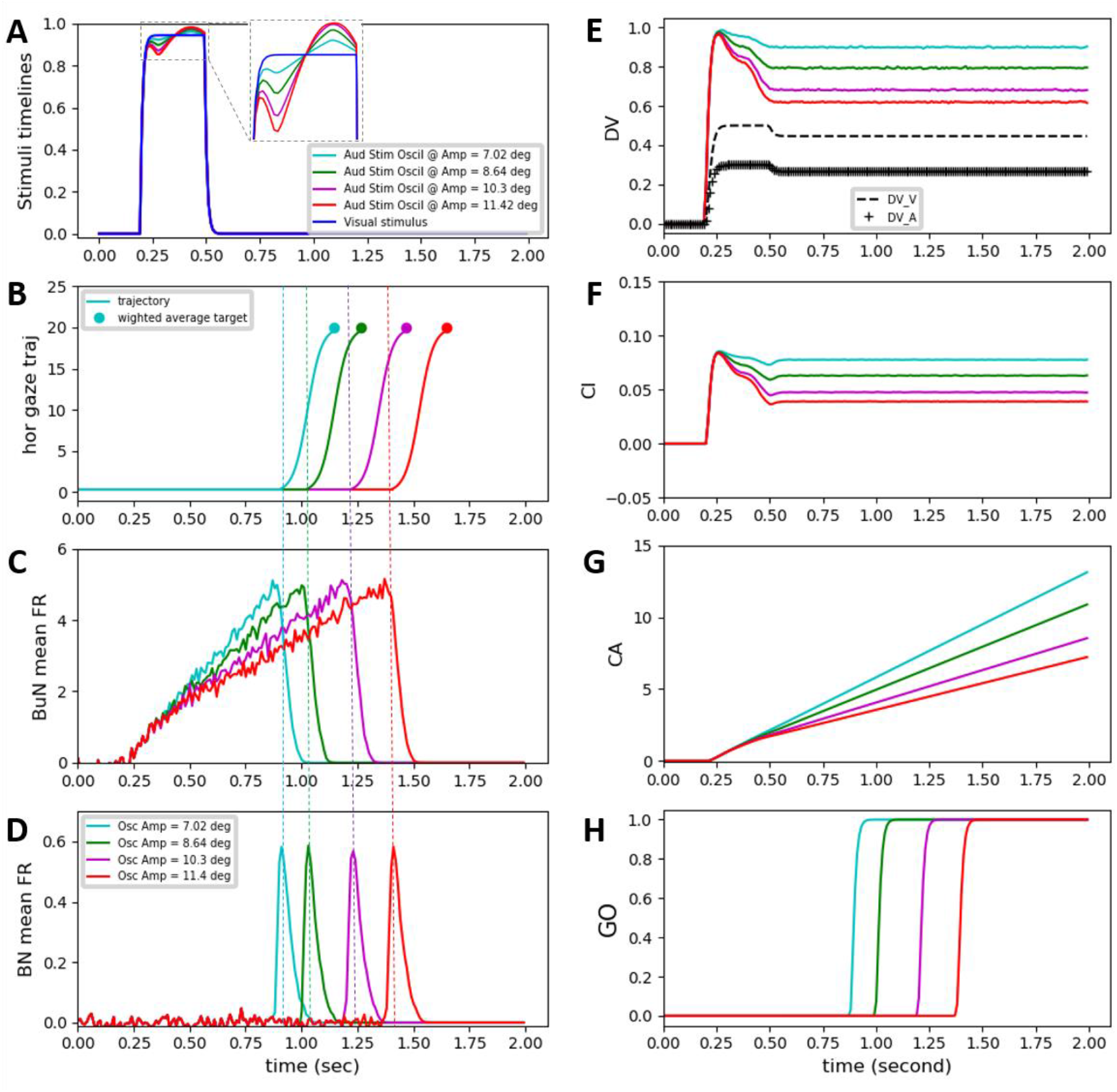
Effect of oscillatory movement of one stimulus on reaction time. Four different conditions have been considered (illustrated by color coding). The visual stimulus has a fixed position, onset time, duration, and reliability for all conditions. The onset time, duration and reliability of the auditory target is also fixed. **A)** The auditory stimulus oscillates with different amplitudes around the stationary visual stimulus. **B)** Gaze position is shown as a function of time. It shows the reaction time in various conditions. **C)** Mean firing rate of build-up neurons in superior colliculus (SC) is shown as a function of time. **D)** Mean firing rate of burst-neurons in SC is shown as a function of time. **E)** The decision variable is shown being developed through time. The unimodal components are the same for all conditions (dashed and crossed lines). The multisensory component is changing between conditions. **F)** The instantaneous confidence on decision is shown through time. The unimodal components are always zero because the multisensory plan is always winning. **G)** The accumulative confidence on decision is shown as a function of time. Only the multisensory component is not zero. **H)** The GO signal is shown for execution of the winning plan. It changes from zero to one whenever the accumulative confidence passes a threshold.

#### Effect of flickering

Another non-tested cross-modal gaze-shift task is when the two stimuli are flickering together. Here we simulate this task (Fig 8), assuming the same onset and flickering frequency for cross-modal stimuli. Both stimuli appear and disappear at the same constant location in space. The spatial distance between the cross-modal stimuli changes in four levels (Fig 8A). RT increases when spatial distance increases (Fig 8B), reflected in the slower build-up of activity in BuNs (Fig 8C) and later response onset of BNs (Fig 8D). This implements the idea that the confidence (Fig 8F, 8G) on the uniqueness of the origin of flickering cross-modal stimuli decreases when their spatial distance increases (Fig 8E), so, the Go signal is issued slower (Fig 8H).

**Figure 8:**
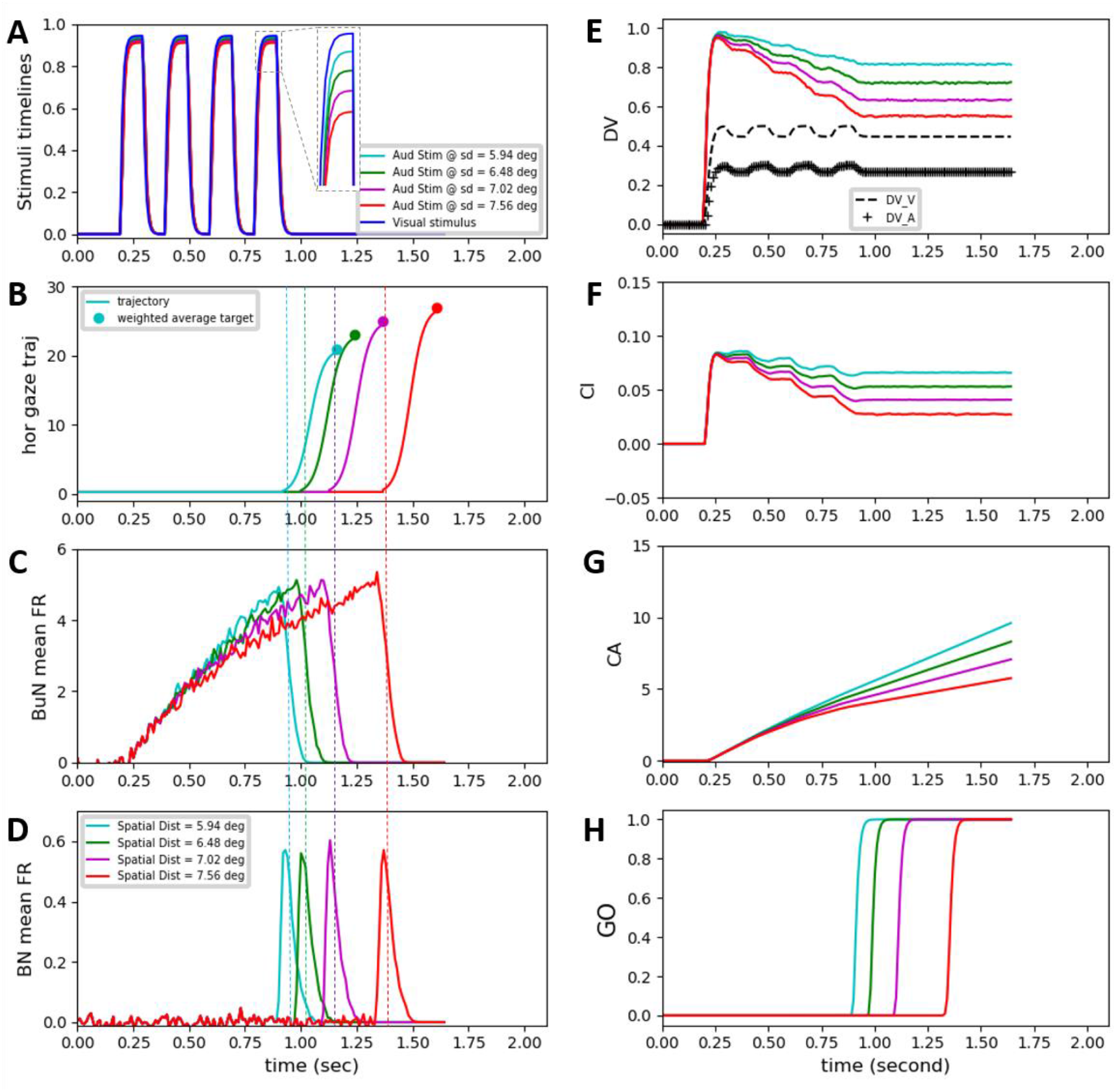
Effect of flickering presentation of cross-modal stimuli on reaction time. Four different conditions have been considered (illustrated by color coding). The visual stimulus has a fixed position, onset time, duration, and reliability for all conditions. The onset time, duration and reliability of the auditory target is also fixed. **A)** The two cross-modal stimuli flicker in time, with four different amounts of spatial distance from each other. **B)** Gaze position is shown as a function of time. It shows the reaction time in various conditions. **C)** Mean firing rate of build-up neurons in superior colliculus (SC) is shown as a function of time. **D)** Mean firing rate of burst-neurons in SC is shown as a function of time. **E)** The decision variable is shown being developed through time. The unimodal components are the same for all conditions (dashed and crossed lines). The multisensory component is changing between conditions. **F)** The instantaneous confidence on decision is shown through time. The unimodal components are always zero because the multisensory plan is always winning. **G)** The accumulative confidence on decision is shown as a function of time. Only the multisensory component is not zero. **H)** The GO signal is shown for execution of the winning plan. It changes from zero to one whenever the accumulative confidence passes a threshold.

### 5.3 Summary of behavioral results

We tested our proposed mechanisms in different tasks. Figure 9 summarizes the preceding results by plotting reaction time as a function of the various task parameters described above. We could replicate these experimentally observed phenomena: 1) the higher the spatial disparity between the stimuli the higher the reaction time, as illustrated in Fig 9A (Frens, Van Opstal, et al., 1995). 2) The higher the temporal disparity between the stimuli the higher the reaction time, as depicted in Fig 9B (Frens, Van Opstal, et al., 1995). 3) If only one of the stimuli is presented, the higher its reliability, the lower the reaction time will be, as explained in Fig 9C (Bell et al., 2006). 4) If two stimuli are presented close to each other in space and time, the higher their reliabilities the higher the reaction time will be, as illustrated in Fig 9D (Diederich & Colonius, 2004). We have also used the model to make predictions in some novel, non-tested, tasks (Fig 9E, 9F, 9G). Our model explains this wide variety of reaction times by introducing various causal structures (represented in alternative plans) for possible environmental phenomena, and suggesting that the relative confidence on a specific causal structure drives the reaction time (see Results for details, and see Discussion for more interpretations).

**Figure 9:**
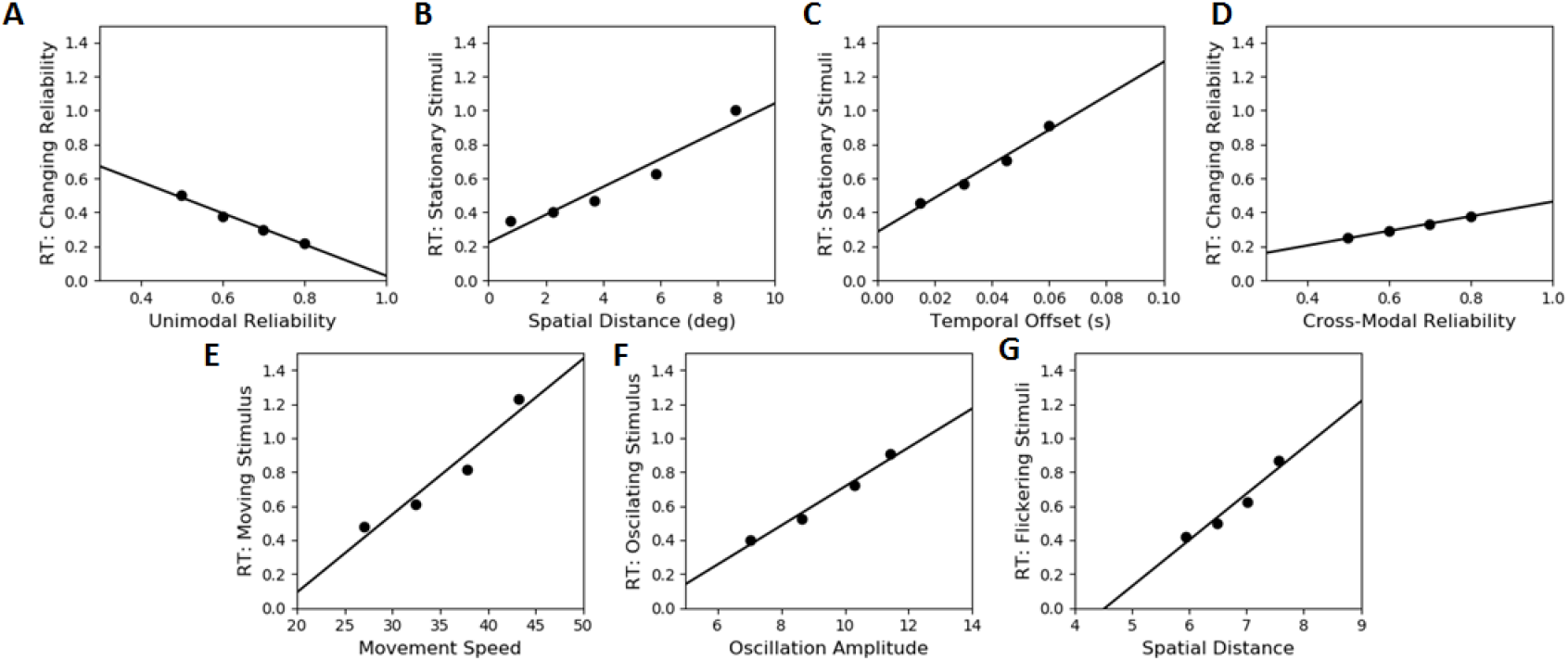
Summary of the behavioral results. This summarizes the predictions of the model for how the reaction time varies as different features of the stimuli change.

We have not concerned ourselves with precise reproduction of quantitative values reported experimentally because of a number of reasons, including: 1) the vast complexity of sensory and perceptual information processing that is affecting the considered behavior but is outside of the scope of our and such similar studies. 2) An exact numerical mapping from the stimulus features manipulated in the experiments onto our intensity and reliability dimensions is not possible. 3) We are keeping our model parameters the same through all our simulations in this and the last paper, while parameters can be changed for each simulation arbitrarily to achieve exact results. Despite all that, we have reproduced all the major trends and even found close numerical results, especially for the more controllable spatial and temporal features.

## 6 Discussion

Here we proposed that the patterns of variability of saccadic reaction times (RT) towards bimodal stimuli are due to high-level cognitive processing. More specifically, the decision-making process for inference of a causal structure, and the confidence on that decision, is proposed to constitute such cognitive processing. Therefore, a high-level conceptual framework (causal inference), implemented at a level that plausibly reflects know physiology, leads to emergent neural signals and behavior that replicates/predicts experimental data

### 6.1 Implications for theories of multisensory action initiation

Previous attempts to model the variability of reaction time towards bimodal stimuli assume the temporal relationships, between the presentations of the two stimuli, as the factor governing the reaction time. Either being race models that consider two separate parallel unimodal channels (Gielen et al., 1983; Raab, 1962), or the coactivation models that consider one additive stage of processing for multimodal stimuli (Diederich, 1992; Schwarz, 1989), or the time-window-of-integration models that combine the two previous ideas (Colonius & Diederich, 2004), they all focus on temporal processing, ignoring the spatial effect (Frens, Van Opstal, et al., 1995). They also isolate this problem from the internal cognitive processing underlying causal inference.

Our model not only considers the effects of both spatial and temporal configuration in a dynamic network, but also relates the perceptual problem of causal inference and the executive problem of action planning in a unifying framework. The model follows the more general idea that when reactionary motor responses towards sensory stimuli are avoided, we allow ourselves to plan actions based on a more complete set of information inferred from the sensory evidence (Chris Eliasmith, 2013). Such inference extends our perception beyond the sensory information, and provides us with a wider range of action plans than sensory-driven reflexive movements (Joaquin M. Fuster, 2005). Selection of one of such action plans and the timing of its execution, then, depends on high-level cognitive processes, rather than reactionary sensorimotor paradigms.

The proposed cognitive system has been characterized by transformations of internal state variables through time. Time evolution of the state space is completely defined, constraining the cognitive system by the limits of planning behaviors in an uncertain, changing environment (Healy & Rowe, 2014). This dynamic nature of the signals in the model provided us with the possibility of designing new psychophysical experiments. We changed the patterns of presentation of the stimuli on the time axis systematically, and saw how the accumulation of evidence, through time, for and against different causal structures, changed the reaction time. Finally, our introduced measure of accumulative confidence can be applied to explain the reaction time in other dynamic decision-making paradigms, where the go command is not forced by the task.

### 6.2 Significance for interpreting previous behavioral findings

It has been observed that the reaction time of planning a gaze-shift towards cross-modal stimuli is affected by the amount of spatial distance between the visual and auditory targets, and by the temporal distance between their presentations (Bell, Meredith, Van Opstal, & Munoz, 2005; Frens, Van Opstal, et al., 1995). These studies rule out statistical facilitation (Raab, 1962) by emphasizing the effect of spatial factor, besides the temporal features, on the facilitation of the reaction time. They hypothesize that the variability of reaction time is due to a multimodal stage of information processing, at a higher level than the primary unisensory processing, although they do not propose a theory for what they hypothesize (Sajad, Sadeh, Yan, Wang, & Crawford, 2016).

The current model accounts for these effects, as illustrated in Figs 3 and 4, by computationally and systematically realizing the intuitive idea that the confidence on the sameness of the origin of the stimuli decreases when the spatial or temporal distance between the stimuli increases. This is accomplished in two steps: 1) introduction of the spatiotemporal similarity of the multimodal stimuli as the criterion for the causal inference, and the saliency of the multisensory plan, 2) defining confidence, driving the reaction time, as how much higher the saliency of the selected plan is than the other alternatives. This meant that the plan to integrate the visual and auditory information and a gaze-shift towards their weighted average becomes less dominant relative to unimodal gaze-shift plans, when the spatial or temporal distance between the stimuli increases. And this leads to a higher reaction time. Thus, our model proposes a cognitive theory for what previous studies hypothesized as a higher-level multisensory stage of processing.

This more general framework enables us to test the system in a wider range of tasks as well. As a first example, in gaze-shifts towards unimodal stimuli, it has been shown that the reaction time decreases by increasing the reliability (intensity) of the stimulus (Bell et al., 2006). They interpreted this as caused by reduced processing time for higher-intensity stimuli, but do not explain why the processing time decreases. Our model explains this phenomenon, as illustrated in Fig 2, by the increased confidence on a unisensory gaze-shift plan, when the stimulus intensity increases. This happens because when, for example, only a visual target is present, the saliencies of the auditory and multisensory plans are zero. So, when the reliability of the visual target increases, the dominance of its corresponding gaze-shift plan increases, and the reaction time decreases.

As another example, in gaze-shifts towards multimodal stimuli presented close to each other in time and space, a reduction in reaction time has been observed when the reliabilities (intensities) of the stimuli change (Diederich & Colonius, 2004). They associate this to the principle of inverse effectiveness in superior colliculus, but do not theorize a mechanism for neither of them. Our model, also, predicts that reaction time increases when the reliabilities (intensities) of the multimodal stimuli increase. This is accounted for by the relative nature of the confidence measure. With a fixed spatiotemporal configuration, and consequently a constant spatiotemporal similarity, the dominance of the multisensory plan relative to the unisensory plans decreases when the saliencies of the unisensory plans, i.e. their stimulus reliabilities, increase. And this leads to a higher reaction time.

### 6.3 Implications for neurophysiology of multisensory processing

Both the causal inference and reaction time parts of the model were designed based on the known neurophysiology about multisensory integration, working memory, decision making, gaze-shift planning and action selection. The functionalities suggested for the expert units, and the transformations realized in their connections, have been defined based on architectural connectivity between different brain areas with known neural behavior. Sustained memory activity, contingent on action, has been shown in posterior parietal cortex (Cohen et al., 1997; J. M. Fuster & Alexander, 1971), in accord with the working memory structures in the causal inference model. The idea of multiple plan representations was inspired by laminar organization of frontal cortex (Berendse et al., 1992; Canteras et al., 1990; Jones et al., 1977; Levesque et al., 1996; Yeterian & Pandya, 1994). The inhibitive effect of decision result on plan representations was considered based on tonic inhibition of cortical and subcortical areas by the basal ganglia (Hikosaka & Wurtz, 1983a, 1983b; Horak & Anderson, 1984). Therefore, implementation of the computational models by an assembly of spiking neural ensembles has explained the multisensory features of SC neurons as emergent properties of the cortical decision-making circuitry (Rohe & Noppeney, 2015; Vivar-Lazo & Fetsch, 2025).

The fundamental evidence that shaped neurophysiological multisensory research was about multimodal neurons in SC. They respond to cross-modal stimuli, aligned in time and space, with a significant enhancement of firing rate compared to their response to the more effective of the unisensory stimuli (Meredith & Stein, 1983; B. E. Stein, Meredith, & Wallace, 1993; Barry E Stein et al., 2020). Our model explains this multisensory enhancement to be caused by the high salience of a plan that represents existence of a common cause for the stimuli, and the high confidence in cortex about the validity of that plan. That is reflected in faster ramp-up of activity in BuNs and earlier burst of activity in BNs. The correspondence between BuN presaccadic buildup with evidence accumulation had been identified in previous studies as well (Ratcliff et al., 2003; Ratcliff et al., 2007).

The SC neurons typically show response enhancement to a spatially aligned multisensory pairing. However, when the stimuli are presented far from each other in space, SC neurons show either no change or a response depression (Frens, Van Opstal, et al., 1995; Meredith & Stein, 1986). The gain of multisensory enhancement reduces by increasing the spatial distance, leading to depression for larger values of spatial distance. Our model explains this trend by the reduction of the confidence in cortex about the notion that the stimuli come from one same source and we should execute a plan towards the location of that source. This can be understood from the slower build-up of activity in BuNs, and later burst of activity in BNs, when the spatial distance increases.

The highest gain of multisensory enhancement of SC neurons’ activity is reported when the stimuli appear at or about the same time. As the temporal misalignment increases, the gain of response enhancement decreases, leading to depression for longer temporal separations (Frens, Van Opstal, et al., 1995; Meredith et al., 1987). This trend is explained by our model through the decline in the confidence on planning a gaze-shift to the weighted average of the two position signals, the inferred common object causing the signals, when the temporal distance between their presentations increases. This is reflected in the slower ramp-up of activity in BuNs, and later burst of activity in BNs, when the temporal distance increases.

Onset latency of multisensory BNs in SC decreases when the intensity of the single-modal stimulus increases (Bell et al., 2006). Our model explains this by the increase in the confidence in cortex on a unimodal plan, which manifests a causal structure of one unimodal source, and making a gaze-shift towards the location of that single source. This can be understood from the faster build-up of activity in BuNs, and earlier burst of activity in BNs, when the unimodal stimulus intensity increases.

In contrast to the unimodal case, the highest gain of multisensory enhancement occurs when the intensities of individual stimuli are weak. As these intensities increase the relative gain of enhancement decreases (Meredith et al., 1987; Porada et al., 2025; B. E. Stein & Stanford, 2008). Our model explains this trend with the decline in the confidence on a unique cause because of the increase in the saliency of the unimodal causal structures, when the intensity of unimodal stimuli increases. This is reflected in the slower ramp-up of activity in BuNs and later burst of activity in BNs when the individual stimulus intensities increase.

### 6.4 Significance for Broader Applications

We have introduced a computational measure of confidence. It is defined within an evidence-based decision-making framework. It is defined on top of the cognitive faculty that gathers and interprets data about the purpose of the task. It consists of an instantaneous measure that denotes how one piece of information informs the decision, and a cumulative measure that denotes how evidence from different pieces of information is gathered together. We showed that this measure could explain the multisensory gaze-shift reaction times. But it can serve to explain other phenomena as well.

Other experiments consisting of the action of gaze-shift towards multisensory stimuli can be also be studied. Their variable timing of action can be explained using our measure of confidence. Some of those experiments have been simulated in the Results: flickering multisensory stimuli, radially moving multisensory stimuli, and vibrating multisensory stimuli. Similar other experiments can be also be designed.

Many other cognitive tasks include planning actions whose initiation times are not forced, but depend on some form of contemplation, resulting in formation of some level of confidence. A modified version of the instantaneous and/or cumulative measures of confidence, defined in this study, can be used to explain whether or not the action is initiated, and also the timing of it. Examples can be found in everyday life actions like when to eat or drink, when to go out, when to pass the street, etc.

This could have applications in the fields of behavioral economics and the so-called neural marketing. Whether or not a consumer buys a product is not an absolutely logical task. The consumer looks at different products. For each product (s)he pays attention to multiple clues, and contemplates whether or not each clue meets his/her desired criterion. If the package designer understands this cognitive process, (s)he can include the critical pieces of information visible on the package, such that the consumer easily, and probably unconsciously, collects those clues and possibly buys that specific product.

### 6.5 Limitations

The model does not explain conditions where more than two stimuli are present. However, we can extend the model to include such scenarios if properly specified.

The model does not explain multisensory conditions where stimuli are of modalities other than visual and auditory, or more than two modalities are present. However, the same model mechanisms can be used to include such conditions as well.

The model does not explain situations where the number of decision alternatives is other than three. However, the evidence-based decision-making architecture can be extended to such conditions.

We are only explaining the behavior of SC neurons in audiovisual saccades. We actually are not claiming that there are specific neurons in SC devoted to this, but we believe they are abstract and robust. The same SC neurons can be recruited for other tasks. Their abstract function is to represent the goal of a gaze-shift. This can be applied to many other tasks.

The model does not include the situations where there are other stimulus dimensions other than spatial disparity, temporal offset, and intensity. Such dimensions could include shape or color for visual stimulus, and linguistic or musical significance for auditory stimulus. More complicated similarity measures should be defined for such situations.

## 7 Conclusion

In this paper, we built up a model of action initiation, on top of our previous causal inference model. The model explained various effects on the reaction time of gaze-shifts towards cross-modal stimuli. The spatial, temporal, and reliability features of the cross-modal stimuli were systematically changed and their effects on the reaction time were reported. In accord with experimental evidence, the reaction time increased when the spatial or temporal distance between the stimuli, or their reliabilities increased.

This model introduced cognitive mechanisms, within the decision-making framework of the previous model, that determine when the winning plan is sent to downstream sensorimotor machinery to be implemented (Daemi & Crawford, 2015; Sparks, 2002). Our model applied the idea of confidence on a winning plan, as the significance of that plan relative to other possible alternative plans, to control the initiation of action. Therefore, we conclude that, in absence of a top-down command to execute the action at a fixed time, the winning plan is executed only when an accumulative measure of confidence, on a selected action plan, reaches a certain threshold.

We then developed the spiking neural network that neurally implements the parallel processing units in the causal inference and reaction time models. This complex network included sub-networks that were expert models of short-term memory in parietal cortex, evidence-based decision making in prefrontal cortex, action selection in basal ganglia, and gaze-shift execution in superior colliculus (SC). SC was the only structure, with clear multisensory experimental evidence, based on which we verified our model. We produced simulations of the behavior of the two SC neural populations in the model, in different multisensory tasks. We showed that the model replicates the multisensory principles of neural behavior in SC, which explained its dependence on the spatial, temporal and reliability properties of the cross-modal stimuli.

We assigned cognitive significance, namely the confidence on choosing a causal structure for the environmental phenomena, to the sensorimotor action planning. This computational measure of confidence can predictably explain other multisensory gaze-shift tasks, and can be generalized to explain other cognitive tasks where the initiation of action is conditioned on collecting enough evidence from the environment in favor of implementing the action. This could have applications in behavioral economics where the consumers’ purchase behavior could be unconsciously guided through presentation of some critical pieces of information about the product.

## 8 Appendix

**Mathematical Formulation**

## 8.1 Decision making

Multimodal signals are detected from the environment and encoded in early sensory areas whose dynamics reflect the temporal aspect of the stimuli presentation. These sensory signals are then communicated to the working memory to be retained and further processed. Spatiotemporal similarity measure, our criterion to infer the origin of the stimuli, is calculated from the sustained signals in the working memory. The plan representations in *PL* are then constructed to manifest the possible causal structures. The two unimodal plans represent the spatial position of the unimodal stimuli along with their reliabilities as the plan saliencies (*sal*_*plv*_ and *sal*_*pla*_). The multisensory plan represents the weighted average of the unimodal position signals along with the spatiotemporal similarity measure as plan saliency (*sal*_*plav*_). The decision variable is constructed by the saliencies of the plans:

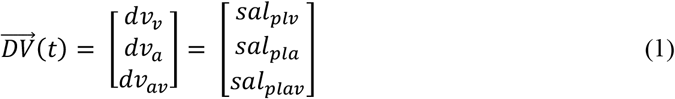

The decision rule is realized by a transformation of the 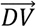 resulting in decision result, 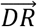:

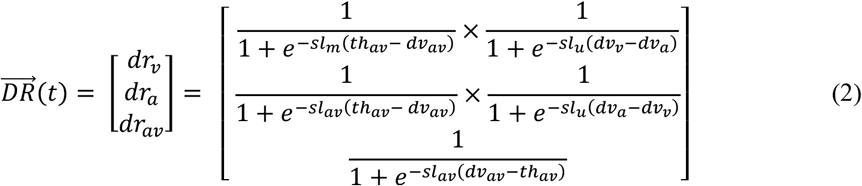

*th*_*av*_ is a threshold value applied on the similarity measure, determining whether or not a unique object originated the signals. 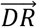 controls the implementation of the decision of which plan to drive the gaze-shift. This is applied by selective inhibition of plan representations in execution, cortical layer (*EX*). Plan representations in *EX* are governed by these equations:

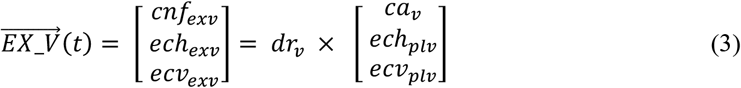

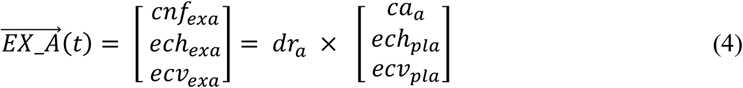

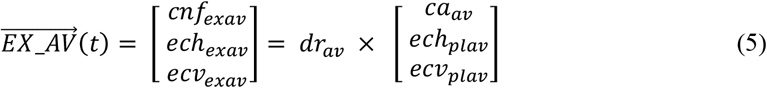

*EX* plan representations are selectively inhibited to determine the winning plan. This effect is shown by the multiplicative effect of the corresponding 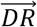 component. We have actually employed a parallel basal ganglia circuitry for implementation of this selection process. However, we are not including the formulations for the basal ganglia computational units because our main focus is on how the decisions are made but not on how they are implemented. *EX* plan representations are assumed to be action initiation confidence maps. The first dimension of *EX* plan representations indicates how confident we are on selecting the corresponding plan if this plan is actually winning. The next section explains how the confidence measures are computed.

## 8.2 Confidence measures

We intend to propose a criterion of when to initiate a gaze-shift when it is free to implement the decision at any time, i.e. the choice-reaction-time case. Conceptually, it is proposed that this timing is determined by the confidence on the decision. First, an instantaneous measure of confidence 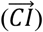 is introduced. The confidence on the decision at each time-point is measured by the distance between the bid of the winning plan and the bids of the losing plans. This variable is constructed as a 3-D signal each component of which indicates at any time, if its corresponding plan is winning, how confident the decision that it is winning is.

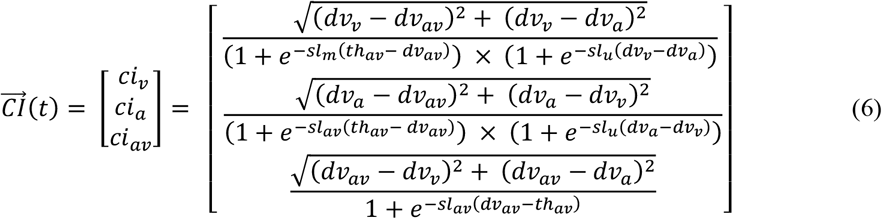

This confidence grows through time with a slope that is dependent on the value of the instantaneous confidence at any time-point. This concept can be materialized in a structure called ‘accumulative confidence’ by integrating the components of ‘instantaneous confidence’ through time. This is a leaky integrator with a controllable leak. The state space vector of this integrator has four dimensions. The first component is the leak and the other three components are the accumulative confidence on the corresponding plans to drive the gaze-shift, if it is winning:

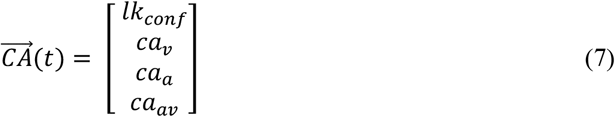

The state space equations characterizing this structure are:

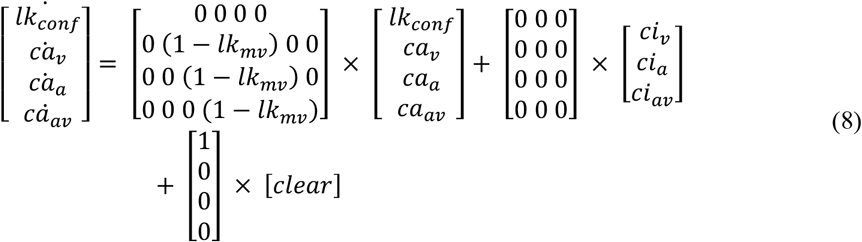

Conceptually, the subject is to make a gaze-shift when its confidence about its decision reaches a threshold.

## 8.3 GO command

We propose that all the plan representations in *EX*, along with their corresponding confidences, converge to the another computational unite that is called 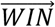:

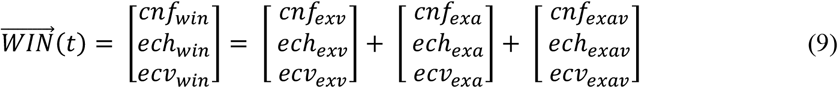

However, because of the selective inhibition applied on *EX* by 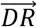, only one plan, which is winning the decision making process, is feeding 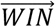 at any time. The confidence map of the winning plan in *EX* is now communicated to 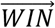. The *GO* command is constructed by transformation of the confidence component of 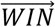:

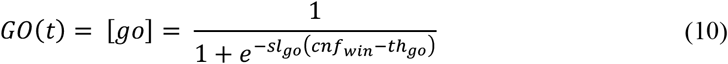

According to this transformation, as soon as the confidence on the winning plan passes a threshold, the value of the *GO* signal changes from 0 to 1. The winning plan is communicated from 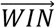 to a final computational unit called 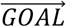.

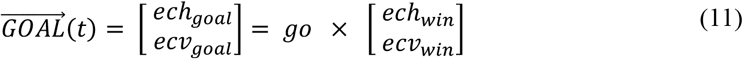

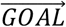 is under constant inhibition of *GO* which determines action initiation. When the *GO* signal has the value zero (the default configuration) the 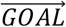 is inhibited. 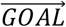 gets released out of the *GO*’s inhibition whenever the *GO* signal becomes one.

### 8.4 Neural Implementation

Cognitive, perceptual, and motor abilities have been associated to the activity of neural circuits in the brain. This mere association has convinced some that cognition will emerge if we can model single neurons’ performance and connect them according to the real synaptic statistics (Markram, 2006). However, cognition has not yet emerged from data-driven large scale models, and there are good reasons to think that cognition may never emerge (C. Eliasmith & Trujillo, 2014). The Neural Engineering Framework (NEF) (Dumont et al., 2023; Kröger, 2023) an approach which starts modeling cognition by describing the system at a higher level of abstraction and then realizing it using neural models with adjustable degrees of accuracy. This method has its roots in the work of a number of researchers (Georgopoulos, Schwartz, & Kettner, 1986; Rieke, 1997; Salinas & Abbott, 1994). The models designed within this framework closely resembles physiological findings in the activity of single neurons, timing of responses, and behavioral errors without being built into the model (Rasmussen & Eliasmith, 2013; Stewart & Eliasmith, 2013). Here we use NEF to assemble local loops in cortex, SC, and BG into a large-scale architecture of connectivity of brain areas interacting for planning a gaze-shift towards cross-modal stimuli. This neurally implements the causal inference (Daemi et al., 2016) and reaction time (first part of this paper) models.

The inputs to the model are transient signals detected from unimodal stimuli in the environment, representing the position and reliability of the sensory stimuli. These transient sensory signals are communicated to and stored in a common short-term memory, the storage part of the working memory. The sensory signals, sustained in short-term memory, are sent to the cognitive-processing part of the working memory where, in our case, a criterion for whether or not to integrate the cross-modal signals is calculated. This criterion captures how close the two stimuli are together by: 1) calculating the spatial distance between them as a function of time, 2) integrating the spatial distance through time. This results in the spatiotemporal similarity measure, which changes between zero and one for the least to most similar stimuli.

We formalize the first part of the cross-modal gaze-shift planning by selecting among three possible causal structures: 1) separate causes for the signals, and target chosen to be the location of the visual stimulus, 2) separate causes for the signals, and target chosen to be the location of the auditory stimulus, 3) unique cause for the signals and target chosen to be the weighted average of the positions of the unimodal stimuli. The three scenarios are realized in three plan representations in the plan layer of FEF. Before making this main decision, we need to determine which of the three scenarios are viable to be considered in the decision making process. This is realized by constructing a 3-D signal from information in the short-term memory, which controls a BG loop to selectively disinhibit the viable plan representations in the FEF plan layer: 1) if no stimuli have been presented, none of the plans are viable, 2) if only one of the stimuli is presented, only the plan corresponding to that modality is viable, 3) if both stimuli are presented, all three plans are viable.

The FEF plan-layer representations are saliency maps of space. Saliency of a unimodal plan is the reliability of its corresponding stimulus, while the saliency of the multisensory plan is the spatiotemporal similarity measure. The saliency of the viable plans are sent from the FEF plan-layer representations to a central decision variable. The spatial maps of the viable plans are sent from the FEF plan-layer to the FEF execution-layer. Choosing between the viable causal structures is materialized by selective disinhibition of the FEF execution-layer representations. A decision rule is realized through the transformation of the decision variable into the decision result: 1) if similarity measure is greater than a threshold, choose the multisensory plan, 2) if the similarity measure is smaller than a threshold and visual stimulus is more reliable than the auditory stimulus, choose the visual plan, 3) if the similarity measure is smaller than a threshold and auditory stimulus is more reliable than the visual stimulus, choose the auditory plan. This decision result controls the BG loop that selectively disinhibits the FEF execution layer representations and materializes the causal inference.

The FEF execution-layer representations are confidence maps of space. Confidence component of these spatial maps represent how confident we are on implementing a plan if that plan is winning the decision making process underlying causal inference. Confidence components are calculated form the decision variable in two steps: 1) first an instantaneous measure of confidence is computed based on the transient distance between the bid of the winning plan relative to the bids of the losing plans, 2) then an accumulative measure of confidence is calculated by integrating the corresponding instantaneous confidence components. Accumulated confidence components are communicated to the FEF execution-layer representations to form the confidence maps of space.

The one disinhibited plan representation in the FEF execution-layer sends its confidence map of space to the SC’s map of space in build-up neurons. Build-up neurons represent the chosen goal of the gaze-shift along with the confidence on execution of the said gaze-shift. Build-up neurons communicate their spatial map to a motor map of space in SC’s burst neurons and its confidence component to form a GO command. GO command is constructed by applying a threshold function on the confidence component, which realizes the idea of controlling the reaction time of the gaze-shift by the accumulated confidence on the inferred causal structure. GO command controls another BG loop that selectively disinhibits SC_B_: constant inhibition until the confidence reaches a threshold. This then sends the gaze-shift motor plan to the brainstem’s eye-head coordination machinery and execute the details of eye and head movements underlying the gaze-shift.

## 9 Acknowledgements

We thank Drs. Jeff Schall and Gunnar Blohm for helpful comments on the manuscript. During this project, Daemi was supported by the CAN-ACT NSERC CREATE Program, and Crawford was supported by the Canada Research Chair Program.

## 10 Nomenclature

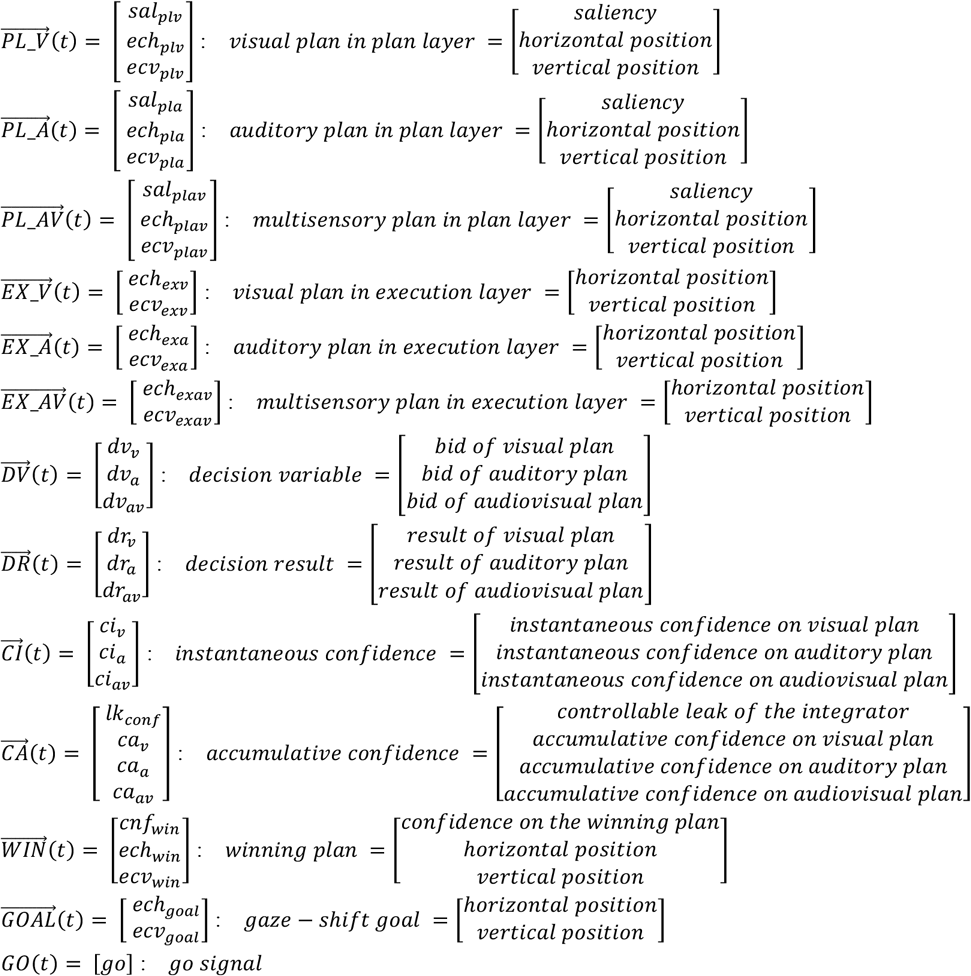

## References

Alvarado, J. C., Stanford, T. R., Vaughan, J. W., & Stein, B. E. (2007). Cortex mediates multisensory but not unisensory integration in superior colliculus. J Neurosci, 27(47), 12775–12786. doi:10.1523/JNEUROSCI.3524-07.2007

Amlot, R., Walker, R., Driver, J., & Spence, C. (2003). Multimodal visual-somatosensory integration in saccade generation. Neuropsychologia, 41(1), 1-15. Retrieved from http://www.ncbi.nlm.nih.gov/pubmed/12427561

Anderson, J. R. (1983). The architecture of cognition. Cambridge, Mass.: Harvard University Press.

Beiser, D. G., & Houk, J. C. (1998). Model of cortical-basal ganglionic processing: encoding the serial order of sensory events. J Neurophysiol, 79(6), 3168–3188. Retrieved from http://www.ncbi.nlm.nih.gov/pubmed/9636117

Bekolay, T., Bergstra, J., Hunsberger, E., Dewolf, T., Stewart, T. C., Rasmussen, D., … Eliasmith, C. (2014). Nengo: a Python tool for building large-scale functional brain models. Front Neuroinform, 7, 48. doi:10.3389/fninf.2013.00048

Bell, A. H., Meredith, M. A., Van Opstal, A. J., & Munoz, D. P. (2005). Crossmodal integration in the primate superior colliculus underlying the preparation and initiation of saccadic eye movements. J Neurophysiol, 93(6), 3659–3673. doi:10.1152/jn.01214.2004

Bell, A. H., Meredith, M. A., Van Opstal, A. J., & Munoz, D. P. (2006). Stimulus intensity modifies saccadic reaction time and visual response latency in the superior colliculus. Exp Brain Res, 174(1), 53–59. doi:10.1007/s00221-006-0420-z

Benedek, G., Mucke, L., Norita, M., Albowitz, B., & Creutzfeldt, O. D. (1988). Anterior ectosylvian visual area (AEV) of the cat: physiological properties. Extrageniculostriate Mechanisms Underlying Visually-Guided Orientation Behavior, 75, 245–255. Retrieved from http://www.ncbi.nlm.nih.gov/pubmed/3187054

Berendse, H. W., Galis-de Graaf, Y., & Groenewegen, H. J. (1992). Topographical organization and relationship with ventral striatal compartments of prefrontal corticostriatal projections in the rat. J Comp Neurol, 316(3), 314–347. doi:10.1002/cne.903160305

Canteras, N. S., Shammah-Lagnado, S. J., Silva, B. A., & Ricardo, J. A. (1990). Afferent connections of the subthalamic nucleus: a combined retrograde and anterograde horseradish peroxidase study in the rat. Brain Res, 513(1), 43–59. doi:0006-8993(90)91087-W [pii]

Cisek, P., & Kalaska, J. F. (2010). Neural mechanisms for interacting with a world full of action choices. Annual Review of Neuroscience, 33, 269–298. doi:10.1146/annurev.neuro.051508.135409

Clarey, J. C., & Irvine, D. R. (1986). Auditory response properties of neurons in the anterior ectosylvian sulcus of the cat. Brain Res, 386(1-2), 12–19. Retrieved from http://www.ncbi.nlm.nih.gov/pubmed/3779403

Cohen, J. D., Perlstein, W. M., Braver, T. S., Nystrom, L. E., Noll, D. C., Jonides, J., & Smith, E. E. (1997). Temporal dynamics of brain activation during a working memory task. Nature, 386(6625), 604–608. doi:10.1038/386604a0

Colonius, H., & Diederich, A. (2004). Multisensory interaction in saccadic reaction time: a time-window-of-integration model. J Cogn Neurosci, 16(6), 1000–1009. doi:10.1162/0898929041502733

Colonius, H., & Diederich, A. (2010). The optimal time window of visual-auditory integration: a reaction time analysis. Front Integr Neurosci, 4, 11. doi:10.3389/fnint.2010.00011

Corneil, B. D., Olivier, E., & Munoz, D. P. (2002). Neck muscle responses to stimulation of monkey superior colliculus. II. Gaze shift initiation and volitional head movements. J Neurophysiol, 88(4), 2000–2018. doi:DOI 10.1152/jn.00960.2001

Corneil, B. D., Van Wanrooij, M., Munoz, D. P., & Van Opstal, A. J. (2002). Auditory-visual interactions subserving goal-directed saccades in a complex scene. J Neurophysiol, 88(1), 438–454. Retrieved from http://www.ncbi.nlm.nih.gov/pubmed/12091566

Daemi, M., & Crawford, J. D. (2015). A kinematic model for 3-D head-free gaze-shifts. Front Comput Neurosci, 9, 72. doi:10.3389/fncom.2015.00072

Daemi, M., Harris, L. R., & Crawford, J. D. (2016). Causal Inference for Cross-Modal Action Selection: A Computational Study in a Decision Making Framework. Front Comput Neurosci, 10, 62. doi:10.3389/fncom.2016.00062

Diederich, A. (1992). Probability inequalities for testing separate activation models of divided attention. Percept Psychophys, 52(6), 714–716. Retrieved from http://www.ncbi.nlm.nih.gov/pubmed/1287576

Diederich, A., & Colonius, H. (2004). Bimodal and trimodal multisensory enhancement: effects of stimulus onset and intensity on reaction time. Percept Psychophys, 66(8), 1388–1404. Retrieved from http://www.ncbi.nlm.nih.gov/pubmed/15813202

Diederich, A., & Colonius, H. (2007). Modeling spatial effects in visual-tactile saccadic reaction time. Percept Psychophys, 69(1), 56–67. Retrieved from http://www.ncbi.nlm.nih.gov/pubmed/17515216

Diederich, A., & Colonius, H. (2008a). Crossmodal interaction in saccadic reaction time: separating multisensory from warning effects in the time window of integration model. Exp Brain Res, 186(1), 1–22. doi:10.1007/s00221-007-1197-4

Diederich, A., & Colonius, H. (2008b). When a high-intensity “distractor” is better then a low-intensity one: modeling the effect of an auditory or tactile nontarget stimulus on visual saccadic reaction time. Brain Res, 1242, 219–230. doi:10.1016/j.brainres.2008.05.081

Dumont, N. S.-Y., Stöckel, A., Furlong, P. M., Bartlett, M., Eliasmith, C., & Stewart, T. C. (2023). Biologically-based computation: How neural details and dynamics are suited for implementing a variety of algorithms. Brain Sciences, 13(2), 245.

Eliasmith, C. (2013). How to build a brain a neural architecture for biological cognition(pp. xvii, 456 pages). Retrieved from http://myaccess.library.utoronto.ca/login?url=http://dx.doi.org/10.1093/acprof:oso/9780199794546.001.0001 http://myaccess.library.utoronto.ca/login?url=http://books.scholarsportal.info/viewdoc.html?id=/ebooks/ebooks2/oso/2014-01-01/1/9780199794546-Eliasmith

Eliasmith, C., Stewart, T. C., Choo, X., Bekolay, T., DeWolf, T., Tang, Y., & Rasmussen, D. (2012). A large-scale model of the functioning brain. Science, 338(6111), 1202–1205. doi:10.1126/science.1225266 338/6111/1202 [pii]

Eliasmith, C., & Trujillo, O. (2014). The use and abuse of large-scale brain models. Curr Opin Neurobiol, 25, 1–6. doi:10.1016/j.conb.2013.09.009 S0959-4388(13)00189-X [pii]

Freedman, E. G., & Sparks, D. L. (1997). Activity of cells in the deeper layers of the superior colliculus of the rhesus monkey: Evidence for a gaze displacement command. J Neurophysiol, 78(3), 1669–1690. Retrieved from <Go to ISI>://A1997XY50300041

Frens, M. A., Hepp, K., Suzuki, Y., & Henn, V. (1996). Rotational kinematics and eye position dependence during vestibular-optokinetic stimulation in the monkey. New Directions in Vestibular Research, 781, 622–624. Retrieved from <Go to ISI>://A1996BF92G00053

Frens, M. A., & Van Opstal, A. J. (1998). Visual-auditory interactions modulate saccade-related activity in monkey superior colliculus. Brain Res Bull, 46(3), 211–224. doi:S0361-9230(98)00007-0 [pii]

Frens, M. A., Van Opstal, A. J., & Van der Willigen, R. F. (1995). Spatial and temporal factors determine auditory-visual interactions in human saccadic eye movements. Percept Psychophys, 57(6), 802–816. Retrieved from http://www.ncbi.nlm.nih.gov/pubmed/7651805

Frens, M. A., Vanopstal, A. J., & Vanderwilligen, R. F. (1995). Spatial and Temporal Factors Determine Auditory-Visual Interactions in Human Saccadic Eye-Movements. Percept Psychophys, 57(6), 802–816. Retrieved from <Go to ISI>://A1995RJ70700006

Fuster, J. M. (2005). Cortex and mind : unifying cognition. Oxford; New York: Oxford University Press.

Fuster, J. M., & Alexander, G. E. (1971). Neuron activity related to short-term memory. Science, 173(3997), 652–654. Retrieved from http://www.ncbi.nlm.nih.gov/pubmed/4998337

Gandhi, N.J., & Katnani, H.A. (2011) Motor functions of the superior colliculus. Annu Rev Neurosci, 34, 205–31. doi: 10.1146/annurev-neuro-061010-113728.

Georgopoulos, A. P., Schwartz, A. B., & Kettner, R. E. (1986). Neuronal population coding of movement direction. Science, 233(4771), 1416–1419. Retrieved from http://www.ncbi.nlm.nih.gov/pubmed/3749885

Gernert, M., Hamann, M., Bennay, M., Loscher, W., & Richter, A. (2000). Deficit of striatal parvalbumin-reactive GABAergic interneurons and decreased basal ganglia output in a genetic rodent model of idiopathic paroxysmal dystonia. J Neurosci, 20(18), 7052–7058. doi:20/18/7052 [pii]

Gielen, S. C., Schmidt, R. A., & Van den Heuvel, P. J. (1983). On the nature of intersensory facilitation of reaction time. Percept Psychophys, 34(2), 161–168. Retrieved from http://www.ncbi.nlm.nih.gov/pubmed/6634374

Girard, B., & Berthoz, A. (2005). From brainstem to cortex: computational models of saccade generation circuitry. Prog Neurobiol, 77(4), 215–251. doi:10.1016/j.pneurobio.2005.11.001

Gold, J. I., & Shadlen, M. N. (2007). The neural basis of decision making. Annual Review of Neuroscience, 30, 535–574. doi:10.1146/annurev.neuro.29.051605.113038

Healy, S. D., & Rowe, C. (2014). Animal cognition in the wild. Behav Processes, 109 Pt B, 101–102. doi:10.1016/j.beproc.2014.11.013

Hershenson, M. (1962). Reaction time as a measure of intersensory facilitation. J Exp Psychol, 63, 289–293. Retrieved from http://www.ncbi.nlm.nih.gov/pubmed/13906889

Hikosaka, O., & Wurtz, R. H. (1983a). Visual and oculomotor functions of monkey substantia nigra pars reticulata. I. Relation of visual and auditory responses to saccades. J Neurophysiol, 49(5), 1230–1253. Retrieved from http://www.ncbi.nlm.nih.gov/pubmed/6864248

Hikosaka, O., & Wurtz, R. H. (1983b). Visual and oculomotor functions of monkey substantia nigra pars reticulata. IV. Relation of substantia nigra to superior colliculus. J Neurophysiol, 49(5), 1285–1301. Retrieved from http://www.ncbi.nlm.nih.gov/pubmed/6306173

Horak, F. B., & Anderson, M. E. (1984). Influence of globus pallidus on arm movements in monkeys. II. Effects of stimulation. J Neurophysiol, 52(2), 305–322. Retrieved from http://www.ncbi.nlm.nih.gov/pubmed/6481435

Jiang, W., Wallace, M. T., Jiang, H., Vaughan, J. W., & Stein, B. E. (2001). Two cortical areas mediate multisensory integration in superior colliculus neurons. J Neurophysiol, 85(2), 506–522. Retrieved from http://www.ncbi.nlm.nih.gov/pubmed/11160489

Jones, E. G., Coulter, J. D., Burton, H., & Porter, R. (1977). Cells of origin and terminal distribution of corticostriatal fibers arising in the sensory-motor cortex of monkeys. J Comp Neurol, 173(1), 53–80. doi:10.1002/cne.901730105

Kiani, R., & Shadlen, M. N. (2009). Representation of confidence associated with a decision by neurons in the parietal cortex. Science, 324(5928), 759–764. doi:10.1126/science.1169405

Klier, E. M., Wang, H., & Crawford, J. D. (2003). Three-dimensional eye-head coordination is implemented downstream from the superior colliculus. J Neurophysiol, 89(5), 2839–2853. Retrieved from http://www.ncbi.nlm.nih.gov/entrez/query.fcgi?cmd=Retrieve&db=PubMed&dopt=Citation&list_uids=12740415

Koos, T., & Tepper, J. M. (1999). Inhibitory control of neostriatal projection neurons by GABAergic interneurons. Nat Neurosci, 2(5), 467–472. doi:10.1038/8138

Kröger, B. J. (2023). The NEF-SPA approach as a framework for developing a neurobiologically inspired spiking neural network model for speech production. Journal of Integrative Neuroscience, 22(5), 124.

Lee, P. H., Helms, M. C., Augustine, G. J., & Hall, W. C. (1997). Role of intrinsic synaptic circuitry in collicular sensorimotor integration. Proc Natl Acad Sci U S A, 94(24), 13299–13304. Retrieved from <Go to ISI>://A1997YJ45600107

Levesque, M., Charara, A., Gagnon, S., Parent, A., & Deschenes, M. (1996). Corticostriatal projections from layer V cells in rat are collaterals of long-range corticofugal axons. Brain Res, 709(2), 311–315. doi:0006-8993(95)01333-4 [pii]

Markram, H. (2006). The blue brain project. Nat Rev Neurosci, 7(2), 153–160. doi:nrn1848 [pii] 10.1038/nrn1848

Meredith, M. A., & Clemo, H. R. (1989). Auditory cortical projection from the anterior ectosylvian sulcus (Field AES) to the superior colliculus in the cat: an anatomical and electrophysiological study. J Comp Neurol, 289(4), 687–707. doi:10.1002/cne.902890412

Meredith, M. A., Nemitz, J. W., & Stein, B. E. (1987). Determinants of multisensory integration in superior colliculus neurons. I. Temporal factors. J Neurosci, 7(10), 3215–3229. Retrieved from http://www.ncbi.nlm.nih.gov/pubmed/3668625

Meredith, M. A., & Ramoa, A. S. (1998). Intrinsic circuitry of the superior colliculus: Pharmacophysiological identification of horizontally oriented inhibitory interneurons. J Neurophysiol, 79(3), 1597–1602. Retrieved from <Go to ISI>://000072525100045

Meredith, M. A., & Stein, B. E. (1983). Interactions among converging sensory inputs in the superior colliculus. Science, 221(4608), 389–391. Retrieved from http://www.ncbi.nlm.nih.gov/pubmed/6867718

Meredith, M. A., & Stein, B. E. (1986). Spatial factors determine the activity of multisensory neurons in cat superior colliculus. Brain Res, 365(2), 350–354. Retrieved from http://www.ncbi.nlm.nih.gov/pubmed/3947999

Mucke, L., Norita, M., Benedek, G., & Creutzfeldt, O. (1982). Physiologic and anatomic investigation of a visual cortical area situated in the ventral bank of the anterior ectosylvian sulcus of the cat. Exp Brain Res, 46(1), 1–11. Retrieved from http://www.ncbi.nlm.nih.gov/pubmed/7067781

Munoz, D. P., Dorris, M. C., & Klein, R. M. (2000). Neural correlates of inhibition of return in the monkey superior colliculus. Perception, 29, 1–1. Retrieved from <Go to ISI>://000207910600003

Munoz, D. P., & Istvan, P. J. (1998). Lateral inhibitory interactions in the intermediate layers of the monkey superior colliculus. J Neurophysiol, 79(3), 1193–1209. Retrieved from <Go to ISI>://000072525100008

Munoz, D. P., & Wurtz, R. H. (1995a). Saccade-Related Activity in Monkey Superior Colliculus. 1. Characteristics of Burst and Buildup Cells. J Neurophysiol, 73(6), 2313–2333. Retrieved from <Go to ISI>://A1995RD79300016

Munoz, D. P., & Wurtz, R. H. (1995b). Saccade-Related Activity in Monkey Superior Colliculus. 2. Spread of Activity during Saccades. J Neurophysiol, 73(6), 2334–2348. Retrieved from <Go to ISI>://A1995RD79300017

Munoz, D. P., & Wurtz, R. H. (1995c). Saccade-related activity in monkey superior colliculus. I. Characteristics of burst and buildup cells. J Neurophysiol, 73(6), 2313–2333. Retrieved from http://www.ncbi.nlm.nih.gov/pubmed/7666141

Munoz, D. P., & Wurtz, R. H. (1995d). Saccade-related activity in monkey superior colliculus. II. Spread of activity during saccades. J Neurophysiol, 73(6), 2334–2348. Retrieved from http://www.ncbi.nlm.nih.gov/pubmed/7666142

Navarra, J., Hartcher-O’Brien, J., Piazza, E., & Spence, C. (2009). Adaptation to audiovisual asynchrony modulates the speeded detection of sound. Proc Natl Acad Sci U S A, 106(23), 9169–9173. doi:10.1073/pnas.0810486106

Navarra, J., Vatakis, A., Zampini, M., Soto-Faraco, S., Humphreys, W., & Spence, C. (2005). Exposure to asynchronous audiovisual speech extends the temporal window for audiovisual integration. Brain Res Cogn Brain Res, 25(2), 499–507. doi:10.1016/j.cogbrainres.2005.07.009

Newell, A., & Simon, H. A. (1972). Human problem solving. Englewood Cliffs, N.J.,: Prentice-Hall.

Porada, D. A., Stein, B. E., & Rowland, B. A. (2025). The Reliability of Multisensory Integration in Enhancing Behavioral Performance. Neuroscience & Biobehavioral Reviews, 106476.

Purcell, B. A., Heitz, R. P., Cohen, J. Y., Schall, J. D., Logan, G. D., & Palmeri, T. J. (2010). Neurally constrained modeling of perceptual decision making. Psychol Rev, 117(4), 1113–1143. doi:10.1037/a0020311

Purcell, B. A., Schall, J. D., Logan, G. D., & Palmeri, T. J. (2012). From salience to saccades: multiple-alternative gated stochastic accumulator model of visual search. J Neurosci, 32(10), 3433–3446. doi:10.1523/JNEUROSCI.4622-11.2012

Raab, D. H. (1962). Statistical facilitation of simple reaction times. Trans N Y Acad Sci, 24, 574–590. Retrieved from http://www.ncbi.nlm.nih.gov/pubmed/14489538

Rasmussen, D., & Eliasmith, C. (2013). Modeling brain function: current developments and future prospects. JAMA Neurol, 70(10), 1325–1329. doi:1729657 [pii] 10.1001/jamaneurol.2013.3835

Ratcliff, R., Cherian, A., & Segraves, M. (2003). A comparison of macaque behavior and superior colliculus neuronal activity to predictions from models of two-choice decisions. J Neurophysiol, 90(3), 1392–1407. doi:10.1152/jn.01049.2002

Ratcliff, R., Hasegawa, Y. T., Hasegawa, R. P., Smith, P. L., & Segraves, M. A. (2007). Dual diffusion model for single-cell recording data from the superior colliculus in a brightness-discrimination task. J Neurophysiol, 97(2), 1756–1774. doi:10.1152/jn.00393.2006

Redgrave, P., Prescott, T. J., & Gurney, K. (1999). The basal ganglia: a vertebrate solution to the selection problem? Neuroscience, 89(4), 1009–1023. doi:S0306452298003194 [pii]

Rieke, F. (1997). Spikes : exploring the neural code. Cambridge, Mass.: MIT Press.

Rohe, T., & Noppeney, U. (2015). Cortical hierarchies perform Bayesian causal inference in multisensory perception. PLoS biology, 13(2), e1002073.

Rubinstein, L. (1964). Intersensory and Intrasensory Effects in Simple Reaction Time. Percept Mot Skills, 18, 159–172. doi:10.2466/pms.1964.18.1.159

Sajad, A., Sadeh, M., Yan, X., Wang, H., & Crawford, J. D. (2016). Transition from Target to Gaze Coding in Primate Frontal Eye Field during Memory Delay and Memory-Motor Transformation. eNeuro, 3(2). doi:10.1523/ENEURO.0040-16.2016

Sajad, A., Sadeh, M., & Crawford, J.D. (2020). Spatiotemporal transformations for gaze control. Physiological Reports. 8 (e14533)

Salinas, E., & Abbott, L. F. (1994). Vector reconstruction from firing rates. J Comput Neurosci, 1(1-2), 89–107. Retrieved from http://www.ncbi.nlm.nih.gov/pubmed/8792227

Schiller, P. H., Sandell, J. H., & Maunsell, J. H. (1987). The effect of frontal eye field and superior colliculus lesions on saccadic latencies in the rhesus monkey. J Neurophysiol, 57(4), 1033–1049. Retrieved from http://www.ncbi.nlm.nih.gov/pubmed/3585453

Schwarz, W. (1989). A new model to explain the redundant-signals effect. Percept Psychophys, 46(5), 498–500. Retrieved from http://www.ncbi.nlm.nih.gov/pubmed/2813037

Sparks, D. L. (2002). The brainstem control of saccadic eye movements. Nat Rev Neurosci, 3(12), 952–964. doi:10.1038/nrn986

Sparks, D. L., & Mays, L. E. (1990). Signal Transformations Required for the Generation of Saccadic Eye-Movements. Annual Review of Neuroscience, 13, 309–336. Retrieved from <Go to ISI>://A1990CT38700017

Stein, B. E., Huneycutt, W. S., & Meredith, M. A. (1988). Neurons and Behavior-the Same Rules of Multisensory Integration Apply. Brain Res, 448(2), 355–358. Retrieved from <Go to ISI>://A1988N415500019

Stein, B. E., Meredith, M. A., & Wallace, M. T. (1993). The visually responsive neuron and beyond: multisensory integration in cat and monkey. Extrageniculostriate Mechanisms Underlying Visually-Guided Orientation Behavior, 95, 79–90. Retrieved from http://www.ncbi.nlm.nih.gov/pubmed/8493355

Stein, B. E., & Stanford, T. R. (2008). Multisensory integration: current issues from the perspective of the single neuron. Nat Rev Neurosci, 9(4), 255–266. doi:10.1038/nrn2331

Stein, B. E., Stanford, T. R., & Rowland, B. A. (2020). Multisensory integration and the society for neuroscience: Then and now. Journal of Neuroscience, 40(1), 3–11.

Sternberg, S. (1969). Memory-scanning: mental processes revealed by reaction-time experiments. Am Sci, 57(4), 421–457. Retrieved from http://www.ncbi.nlm.nih.gov/pubmed/5360276

Stewart, T. C., & Eliasmith, C. (2013). Realistic neurons can compute the operations needed by quantum probability theory and other vector symbolic architectures. Behav Brain Sci, 36(3), 307–308. doi:10.1017/S0140525X12003111S0140525X12003111 [pii]

van Gelder, T. (1998). The dynamical hypothesis in cognitive science. Behav Brain Sci, 21(5), 615-628; discussion 629-665. Retrieved from http://www.ncbi.nlm.nih.gov/pubmed/10097022

Van Wanrooij, M. M., Bremen, P., & John Van Opstal, A. (2010). Acquired prior knowledge modulates audiovisual integration. Eur J Neurosci, 31(10), 1763–1771. doi:10.1111/j.1460-9568.2010.07198.x

VanOpstal, A. J., & Frens, M. A. (1996). Task-dependence of saccade-related activity in monkey superior solliculus: Implications for models of the saccadic system. Extrageniculostriate Mechanisms Underlying Visually-Guided Orientation Behavior, 112, 179–194. Retrieved from <Go to ISI>://A1996BJ08P00013

Vivar-Lazo, M., & Fetsch, C. R. (2025). Neural basis of concurrent deliberation toward a choice and confidence judgment. Nature neuroscience, 1–12.

Wallace, M. T., Meredith, M. A., & Stein, B. E. (1993). Converging influences from visual, auditory, and somatosensory cortices onto output neurons of the superior colliculus. J Neurophysiol, 69(6), 1797–1809. Retrieved from http://www.ncbi.nlm.nih.gov/pubmed/8350124

Wurtz, R. H., & Goldberg, M. E. (1972). The role of the superior colliculus in visually-evoked eye movements. Bibl Ophthalmol, 82, 149–158. Retrieved from http://www.ncbi.nlm.nih.gov/pubmed/4631287

Yeterian, E. H., & Pandya, D. N. (1994). Laminar origin of striatal and thalamic projections of the prefrontal cortex in rhesus monkeys. Exp Brain Res, 99(3), 383–398. Retrieved from http://www.ncbi.nlm.nih.gov/pubmed/7957718

